# BWR-finder: BWT-based *de novo* interspersed repeat detection for gigantic genomes

**DOI:** 10.64898/2026.08.03.738468

**Authors:** Atsushi Takeda, Tsukasa Fukunaga, Michiaki Hamada

## Abstract

We developed BWR-finder (Burrows–Wheeler transform-based Repeat finder), a new software tool for database-free detection of interspersed repeats in large genomes of tens of gigabases. BWR-finder employs a BWT-based seed-and-extend repeat detection algorithm and parallelized extension computation, improving both runtime and memory usage. In benchmarks using the rice and human genomes, BWR-finder reduced runtime compared with RepeatModeler2, HiTE, and REPrise, and reduced memory usage compared with HiTE and REPrise, while maintaining high nucleotide-level repeat detection sensitivity. BWR-finder also enabled whole-genome repeat detection in genomes larger than 10 Gb and identified candidate repeat regions and repeat consensus sequences in the 20.3-Gb *Pleurodeles waltl* genome that were not associated with existing repeat annotations or libraries.

## 1 Introduction

Interspersed repeats are subsequences that occur repeatedly throughout genomes and constitute a large fraction of many eukaryotic genomes. Many interspersed repeats are derived from transposable elements (TEs). The composition of TEs varies substantially among species [1]. This variation reflects the lineage-specific amplification of different TEs during genome evolution. Therefore, TE analysis needs to be performed for each species to annotate TE-derived regions accurately [2].

The proportion of interspersed repeats in a genome tends to increase with genome size [3]. For example, approximately 85% of the 14.6-Gb wheat(*Triticum aestivum*) genome [4] and approximately 90% of the 34.6 Gb lungfish(*Neoceratodus forsteri* ) genome are derived from TEs [5]. Comparative analyses among closely related lineages suggest that the self-proliferation of TEs has contributed to host genome expansion [6]. Thus, TE annotation in large genomes is essential for understanding the mechanisms underlying genome expansion.

Interspersed repeats are typically annotated in genomes by aligning repeat libraries, such as Dfam [7] and Repbase [8], to genomic sequences using RepeatMasker [9]. In this process, expert-curated repeat libraries are commonly used. Because TEs are highly diverse among species, repeat libraries need to be constructed for indiviual species. However, the rapid increase in newly sequenced genomes has outpaced the development of curated repeat libraries for many non-model species. Therefore, methods have been developed to construct repeat libraries directly from genomic sequences without expert manual curation. Methods for constructing interspersed repeat libraries from genomic sequences can be broadly classified into two categories: structure-based methods and *de novo* methods. Structure-based methods detect repeats by using structural features conserved in TEs. For example, LTR retriever [10] uses LTR structures, whereas HelitronScanner [11] detects sequence structures specific to rolling-circle TEs. These methods achieve high accuracy for specific TE classes. However, they do not cover all TE classes, and their ability to detect ancient TEs is limited when characteristic structures have been lost through mutation or fragmentation.

In contrast, *de novo* methods identify repetitive regions based on sequence similarity within a genome. From the perspective of algorithm design, these methods can be divided into seed-and-extend approaches and self-alignment approaches. Seed-and-extend approaches use frequently occurring short sequences as seeds and construct consensus sequences by extending alignments in both directions. Representative examples include RepeatScout [12] and REPrise [13]. Self-alignment approaches perform genome-wide pairwise alignments and identify interspersed repeats by grouping similar regions. Representative examples include RECON [14] and HiTE-FMEA, the *de novo* TE-searching module of HiTE [15]. Because seed-and-extend approaches use greedy multiple sequence alignment, they have an advantage in computational speed. In contrast, self-alignment approaches are based on pairwise alignments, which are generally more accurate than multiple sequence alignment, and are therefore better suited for detecting highly diverged repeats. RepeatModeler2 [16], one of the most widely used repeat detection software packages, achieves sensitive repeat detection by combining both strategies in its *de novo* module.

Although many tools have been developed, existing *de novo* methods for detecting interspersed repeats remain difficult to apply to large genomes because of their computational requirements. Existing seed-and-extend methods require not only the genome sequence but also additional data structures, such as suffix arrays or hash tables, to access seed occurrences. These data structures lead to high memory consumption. In the analysis of the human genome, which is approximately 3.1 Gb, REPrise [13] requires approximately 37 GB of memory even in its most memory-efficient mode. As genome size increases further, this memory requirement exceeds a practical level. Self-alignment approaches can impose a substantial computational burden on long genomes because they require searching for and aligning many pairs of similar genomic regions. HiTE [15] mitigates this burden by partitioning the genome assembly into smaller chunks and masking tandem repeats(TRs) before applying its *de novo* TE-searching procedure. Nevertheless, for the approximately 2.2-Gb *Zea mays* genome, HiTE required 12 GB of memory and 25 h 44 min of runtime. RepeatModeler2 [16] reduces computational cost by sampling the input genome and using only 400 Mb of the genome for repeat detection. However, this strategy cannot detect interspersed repeats located in unsampled regions. Thus, methods that can analyze entire large genomes as input while keeping computational requirements low have not yet been sufficiently established.

The Burrows–Wheeler transform (BWT) [17] is a sequence transformation technique widely used for handling large sequences. The BWT is a reversible transformation based on the suffix array and serves as the basis of the Ferragina–Manzini index (FM-index) [18], a full-text index that enables sequence search in a compressed representation. In addition, by storing the BWT sequence using succinct data structures such as wavelet trees, neighboring substrings around arbitrary positions can be restored without retaining the original sequence. Because of these properties, the BWT has been widely used as a core technology in read-mapping tools [19–21].

Here, we propose BWR-finder, an interspersed repeat detection software tool that uses the BWT to enable analysis of large-scale genomes. BWR-finder adopts an algorithm that performs seed-and-extend-based detection on the BWT sequence. By implementing seed search and each operation in the extension step as BWT-based operations using an FM-index data structure, BWR-finder detects repeats with low memory usage without retaining the original sequence or additional large-scale data structures. Furthermore, BWR-finder accelerates repeat detection through parallel extension, based on the fact that each seed can be processed independently during the extension step. This approach enables interspersed repeat detection from whole large-genome inputs using practical computational resources.

## 2 Results

### 2.1 Overview of BWR-finder

BWR-finder takes an assembled genome sequence or a TR-masked genome sequence as input and outputs a repeat library, which is a set of consensus sequences representing detected repeats. The workflow is shown in Fig.1(A). The input sequence is converted into a BWT sequence using grlBWT [22], which can construct the BWT with low memory usage for large and highly repetitive sequences. The BWT sequence is then stored in memory using the wavelet tree library implemented in sdsl-lite [23]. The main algorithm of BWR-finder is based on the seed-and-extend framework. BWR-finder performs the extension step in parallel using multiple threads, based on the fact that extension after seed search can be processed independently for each seed. Finally, CD-HIT [24] is applied to reduce redundancy in the detected repeat library.

**Fig. 1.**
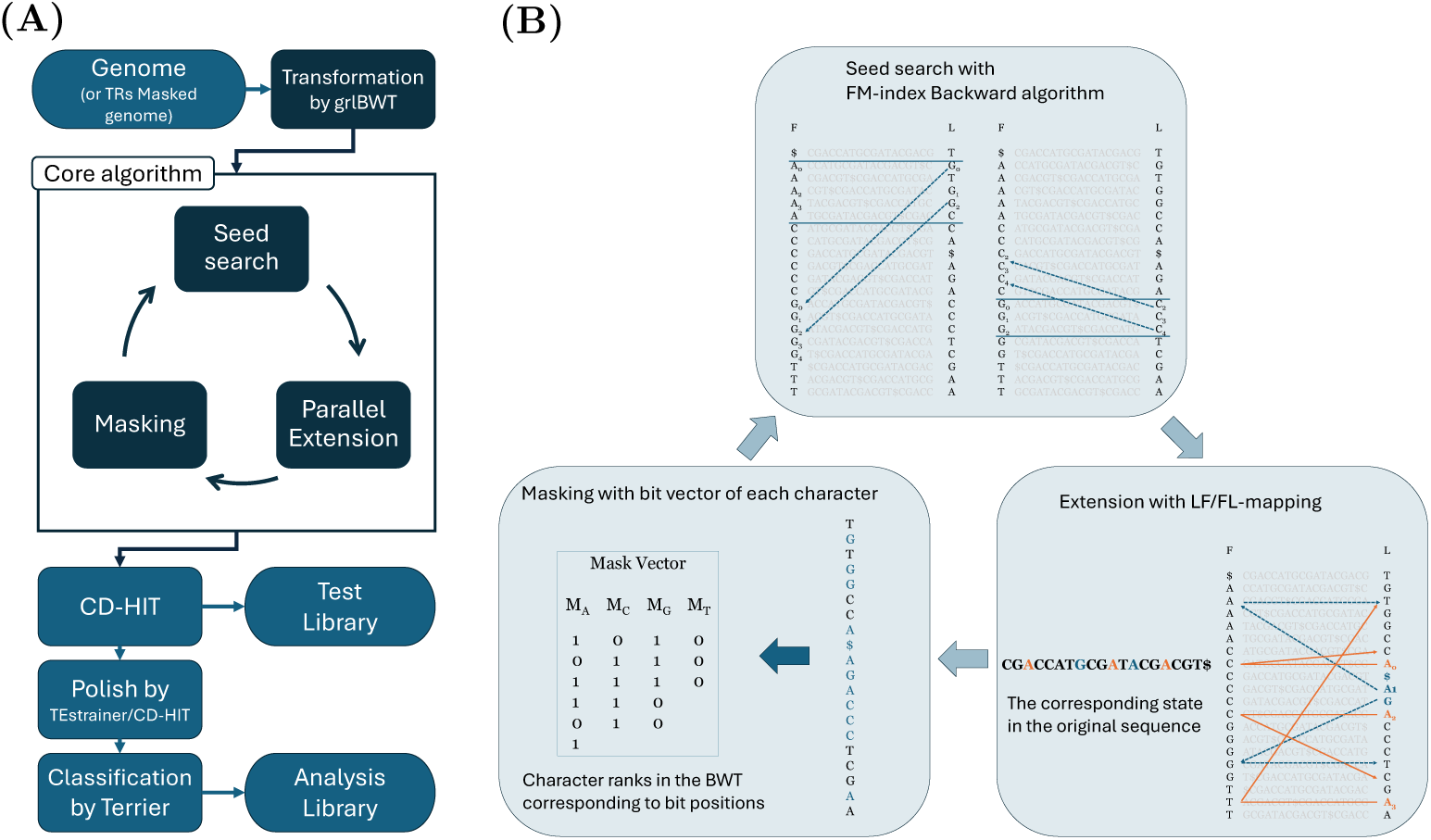
(**A**) Overview of the proposed algorithm. The Test Library is used in Results 2.4–2.6, whereas the Analysis Library is used in Results 2.7. **(B)** Detailed overview of the core algorithm of BWR-finder. In the seed search step, seeds are identified using an FM-index backward-search-like procedure. In the extension step, candidate sequences are extended outward from the seed positions detected in the seed search step using LF/FL mapping. In the masking step, separate bit vectors are prepared for the four nucleotide characters A, C, G, and T, and each position is evaluated based on the rank of the corresponding character in the BWT sequence.

#### 2.1.1 Interspersed repeat detection using a BWT-based seed-and-extend algorithm

BWR-finder performs seed-and-extend-based interspersed repeat detection on the BWT representation of the genome (Fig. 1(B)). This representation supports both locating substring occurrences and accessing neighboring characters in the original sequence. In the seed step, the occurrences of each seed sequence are located. For this purpose, RepeatScout uses a hash table that stores *k*-mer occurrences, whereas REPrise uses a suffix array as an additional data structure for seed indexing. In contrast, BWR-finder performs FM-index-based seed search directly on the BWT.

In the extension step, BWR-finder evaluates sequence similarity around each seed while extending the surrounding regions one base at a time. RepeatScout and REPrise perform this process by updating positions on the original sequence, namely by applying *position* ± 1. In contrast, BWR-finder performs extension alignment by applying LF/FL mapping, which is an operation for searching adjacent characters on the BWT sequence. Thus, BWR-finder completes both seed search and the extension step on the BWT sequence and does not retain additional large-scale data structures.

We next compare the memory usage of these methods. Let *n* be the sequence length, *m* be the total number of seeds, and *σ* be the alphabet size. For DNA sequences, *σ*is at most approximately 6, even when symbols representing masked regions and terminal symbols are included in addition to the four bases A, C, G, and T. RepeatScout theoretically requires *O*(*n* log *σ*+*m* log *n*) bits of memory because it retains the original sequence and a k-mer occurrence hash table. REPrise requires *O*(*n* log *σ* + *n* log *n*) bits of memory because it retains the original sequence and the suffix array. However, BWR-finder stores the BWT sequence in a wavelet tree, a compact data structure that enables efficient search and sequence traversal ( Supplementary Figure S2). In the sdsl-lite implementation [23], the BWT-wavelet tree can be stored in approximately *O*(*αn* log *σ*) bits, where *α* ≒ 1.25, making it more memory-efficient than the two conventional methods. Details are provided in Methods.

#### 2.1.2 Parallel extension step

In repeat detection, most of the computational time is spent in the extension step. BWR-finder parallelizes the extension step across threads by exploiting the fact that the extension process for each seed is independent of those for other seeds. This parallelization reduces the computational time required for the extension step.

However, parallelizing the extension step raises an important design issue: whether and how to control the processing order of seeds. REPrise and RepeatScout preferentially extend high-frequency seeds. This processing order allows these methods to process seeds that represent major repeat consensus sequences earlier. As a result, other seeds derived from the same repeat consensus sequence are less likely to be extended as independent repeat candidate sequences.

In contrast, BWR-finder does not control the extension order based on seed frequency in order to facilitate parallel execution. This design may increase redundant sequences derived from the same repeat consensus sequence, as well as fragmented sequences that reflect only part of a consensus sequence.

Redundant sequences can potentially be consolidated by downstream clustering with CD-HIT [24]. In contrast, CD-HIT has limited ability to reconstruct a complete sequence from fragmented candidates. Therefore, although BWR-finder reduces computational time by parallelizing the extension step, this design may increase the number of fragmented sequences. This trade-off between computational efficiency and sequence completeness may affect the quality of the final repeat library.

### 2.2 Downstream processing

In this study, we constructed two types of libraries and used them for different purposes: evaluating the repeat detection performance of BWR-finder itself and analyzing repeats in *Pleurodeles waltl* (Fig. 1). For the evaluation of repeat detection performance, we constructed a library by applying only CD-HIT to the output of BWR-finder; this library is referred to as the Test Library. For repeat analysis, we constructed another library that included additional refinement and classification steps, because practical repeat annotation generally requires refinement of consensus sequences and classification of repeat types. Specifically, after running CD-HIT, we used TEstrainer, the polishing module included in the repeat detection pipeline Earl-Grey [25], for refinement, and Terrier [26] for sequence classification. This library is referred to as the Analysis Library.

### 2.3 Evaluation metrics

To evaluate the detection performance of BWR-finder and the quality of the resulting repeat libraries, we used three evaluation metrics adopted in a previous study [15](Fig.2). In this study, we refer to these metrics as BM EDTA, BM HiTE, and BM RM2.

**Fig. 2.**
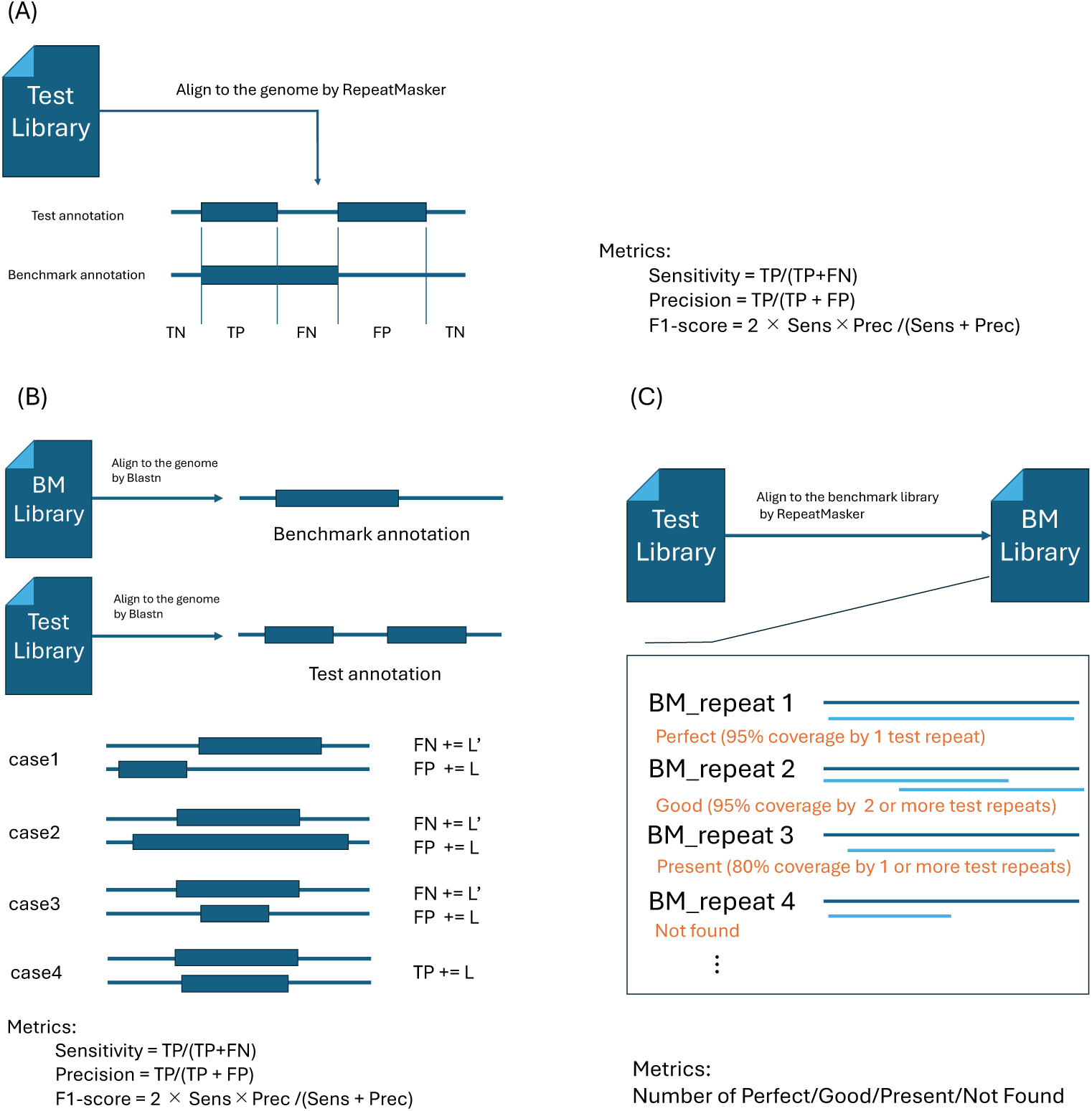
Three benchmark metrics. TP = true positive, TN = true negative, FP = false positive, FN = false negative. BM = Benchmark.(A) BM EDTA. Benchmark annotations and test annotations are compared, and each nucleotide position is classified as one of the binary evaluation outcomes. (B) BM HiTE. For each pair of overlapping annotations, *L* and *L^′^* denote the lengths of the Test annotation and the benchmark annotation, respectively. Test annotations and benchmark annotations are generated by BLASTN searches of repeat libraries against the genome. The coverage between the test and benchmark annotations is then evaluated, and TP is counted only when the test and benchmark annotations overlap reciprocally by at least 80%, as illustrated in case 4. (C) BM RM2. The Test Library and benchmark library are aligned using RepeatMasker. Repeats in the benchmark library are classified into four categories, Perfect, Good, Present, and Not found, based on alignment coverage.

BM EDTA (Fig.2(A)) is an evaluation framework that compares predicted repeat annotations with benchmark annotations based on nucleotide-level coverage. Because this metric directly compares predicted repeat annotations with exisiting repeat annotations, it has been widely used in evaluations of *de novo* repeat detection methods [27, 28].

BM HiTE (Fig.2(B)) evaluates whether repeats are detected as full-length sequences relative to the benchmark annotation. In this framework, the detected repeat consensus sequences are re-annotated in the genome using BLASTN [29], and their coverage is compared with that of the benchmark annotation. Up to this point, BM HiTE is similar to BM EDTA. However, in BM HiTE, True Positive(TP) is counted only when the benchmark annotation and the test annotation overlap reciprocally by at least 80%. In other words, the overlap must cover at least 80% of both the benchmark annotation and the test annotation, thereby limiting excess or deficient predictions. In addition, redundancy or fragmentation in the test annotation increases the False Positive(FP) value.

Unlike the two annotation-based metrics described above, BM RM2 (Fig. 2(C)) directly evaluates how well the constructed repeat library recovers the benchmark repeat library. The predicted repeat library and the benchmark repeat library are aligned using RepeatMasker, and repeat sequences in the benchmark library are classified into four categories: “Perfect” (95% coverage by one test repeat), “Good” (95% coverage by two or more test repeats), “Present” (80% coverage by one or more test repeats), and “Not found”.

Using runtime, memory usage, and the three metrics described above, we compared BWR-finder, RepeatModeler2, HiTE, and REPrise on the rice [30] and human genomes [31, 32]. Because BWR-finder detects repetitive sequences based on their sequence recurrence and does not explicitly distinguish TRs from interspersed repeats, we evaluated BWR-finder both with and without prior masking of TRs by tantan[33]. We also evaluated runtime and memory usage for BWR-finder and RepeatModeler2 on large genomes: wheat (*Triticum aestivum*; 14.6 Gb)[34, 35],salamander (*Pleurodeles waltl* ; 20.3 Gb) [36], and two lungfish species (*Neoceratodus forsteri* [5]; 34.6 Gb, and *Lepidosiren paradoxa* [37]; 87.2 Gb). The other methods were not included in this large-genome benchmark because of their runtime and memory requirements.

Details of the genome assemblies and annotations used in these analyses are provided in Supplementary Table S1.

We evaluated detection performance using BM EDTA and BM RM2 only for the wheat and salamander genomes. Because large genomes may contain repeat sequences that are absent from existing repeat annotations, the BM EDTA-based comparison used %Positive, defined as the proportion of the genome annotated as repeats, and Potential Novel Repeat (PNR), defined as the proportion of the genome classified as FP, in addition to sensitivity. Detection performance evaluation for the lungfish genomes and BM HiTE evaluation for large genomes were not performed because of computational time constraints.

All experiments were performed on a compute node equipped with AMD EPYC 9654 processors (96 cores, 2.4 GHz). Each tool was run with 32 parallel threads and a total allocated memory of 256 GB. Detailed information on software versions and representative commands is provided in Supplementary Table S2 and Supplementary Method S1.6.

### 2.4 Benchmarking on the Rice genome(382 Mb)

We compared BWR-finder (TRs masked and TRs not masked), RepeatModeler2, HiTE, and REPrise using the rice genome (*Oryza sativa*; 382 Mb) (Fig. 3).

**Fig. 3.**
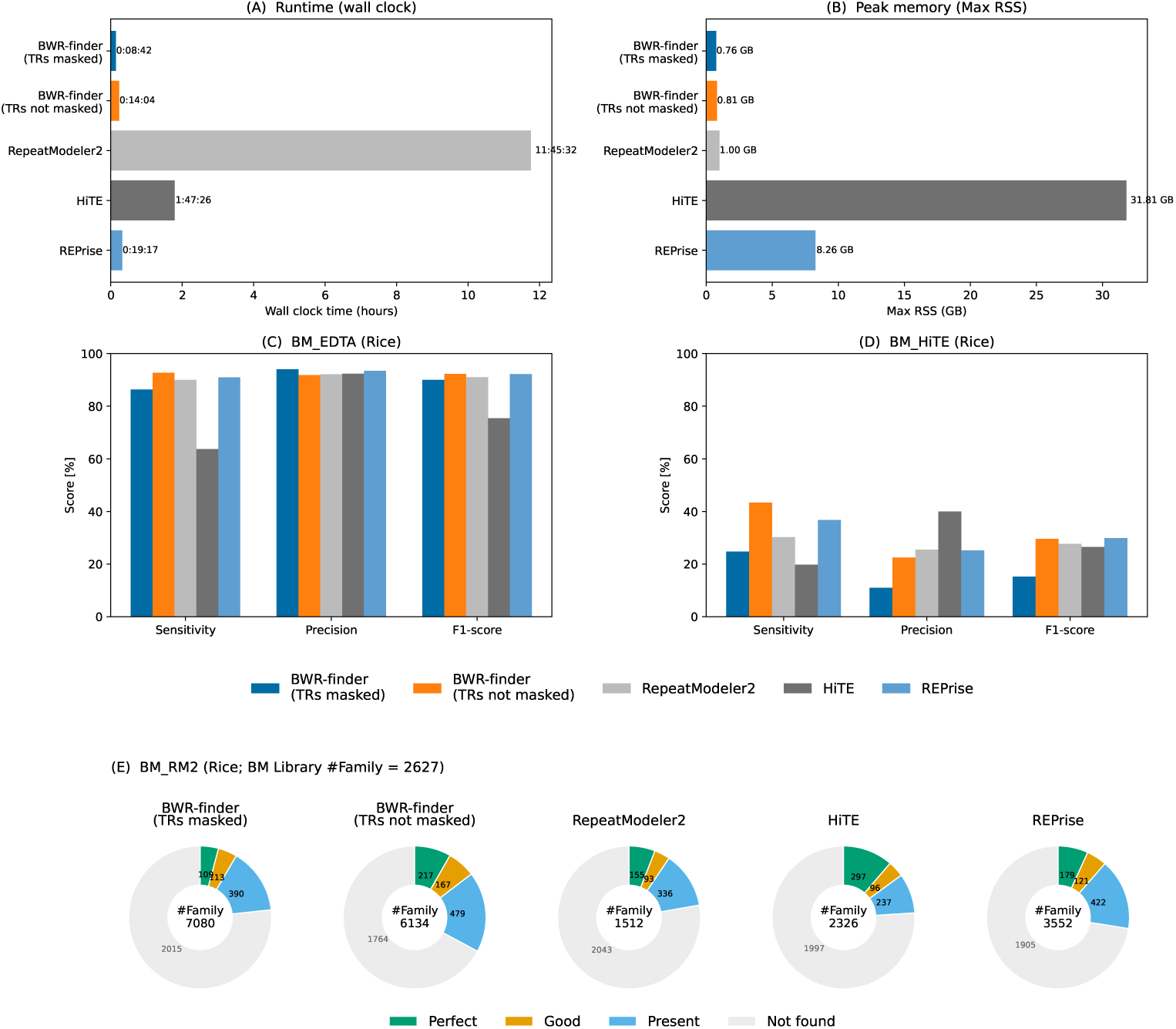
Benchmark results of BWR-finder on the rice genome. (A) Runtime. (B) Memory usage. (C) BM EDTA. (D) BM HiTE. (E) BM RM2. The numbers inside the donut holes indicate the number of consensus sequences in each Test Library.

BWR-finder (TRs masked) completed the analysis in 8 min 42 s. This was approximately 80 times faster than RepeatModeler2 (approximately 11 h 46 min), approximately 12 times faster than HiTE (approximately 1 h 47 min), and approximately two times faster than REPrise (19 min 17 s). BWR-finder (TRs not masked) also completed the analysis in 14 min, indicating that TR masking had only a limited effect on runtime (Fig. 3(A)). The maximum memory usage of BWR-finder was the lowest among the tested methods, ranging from 0.76 to 0.81 GB, and was comparable to that of RepeatModeler2 (1.00 GB). In contrast, REPrise and HiTE required 8.26 GB and 31.81 GB of memory, respectively (Fig. 3(B)). These results indicate that BWR-finder was the most computationally efficient method for the rice genome.

In the nucleotide-level evaluation using BM EDTA (Fig. 3(C)), BWR-finder (TRs not masked) achieved the highest sensitivity, at 92.71%. This sensitivity was 1–3 % higher than those of REPrise (90.96%) and RepeatModeler2 (89.96%) and substantially higher than that of HiTE (63.74%). Precision ranged from 91% to 95% across all methods, with no large differences among methods. These results indicate that BWR-finder achieved detection performance comparable to or higher than that of existing methods. TR masking reduced sensitivity by 6.37 percentage points, suggesting that some interspersed repeats were also masked during TR masking. In contrast, precision increased by 2.16 %, and consequently, the decrease in F1-score was limited to 2.26 %. In BM HiTE (Fig. 3(D)), BWR-finder (TRs not masked) achieved the highest sensitivity, at 43.34%, followed by REPrise (36.75%), RepeatModeler2 (30.24%), BWR-finder (TRs masked) (24.73%), and HiTE (19.79%). In contrast, HiTE achieved the highest precision, at 40.01%, followed by RepeatModeler2 (25.47%), REPrise (25.21%), and BWR-finder (22.47%). BM HiTE is a metric in which fragmented repeats can increase FP counts redundantly for the same repeat region, thereby substantially decreasing precision. HiTE had the smallest total number of bases detected as full-length repeats (approximately 12 Mb), but it also had the smallest total number of bases detected as non-full-length repeats (approximately 18 Mb). In contrast,

BWR-finder and REPrise had larger numbers of bases detected as non-full-length repeats: 123 Mb for BWR-finder (TRs masked), 93 Mb for BWR-finder (TRs not masked), and 69 Mb for REPrise (Supplementary Table S5). These results indicate that although REPrise and BWR-finder have high detection sensitivity, their outputs contain many fragmented repeat consensus sequences.

In the library-sequence-level evaluation using BM RM2 (Fig. 3(E)), HiTE produced the largest number of Perfect models (297), followed by BWR-finder (TRs not masked) (217), REPrise (179), RepeatModeler2 (155), and BWR-finder (TRs masked) (109). In contrast, the cumulative number of sequences classified as Perfect, Good, or Present was highest for BWR-finder (TRs not masked) (863), followed by REPrise (722), HiTE (630), and RepeatModeler2 (584). The cumulative number for BWR-finder (TRs masked) was 612, indicating that TR masking reduced library recovery, consistent with the BM EDTA results. In addition, compared with the 2,627 sequences in the exisiting repeat library, BWR-finder produced 7,080 sequences in the TRs masked condition and 6,134 sequences in the TRs not masked condition. These numbers were larger than those obtained by RepeatModeler2, HiTE, and REPrise (1,512, 2,326, and 3,552 sequences, respectively). Thus, the BM RM2 results also indicate that BWR-finder achieved the highest recovery of the benchmark repeat library, while producing a more fragmented set of detected repeat consensus sequences.

### 2.5 Benchmarking on the Human genome(3.1Gb)

Similarly, we compared the computational requirements and detection performance of BWR-finder (TRs masked and TRs not masked), RepeatModeler2, HiTE, and REPrise using the human genome (Fig. 4). BWR-finder (TRs masked) completed the analysis in 1 h 50 min. This was approximately 15 times faster than HiTE (approximately 28 h), approximately 5.5 times faster than RepeatModeler2 (approximately 10 h), and more than two times faster than REPrise (approximately 4.5 h). BWR-finder (TRs not masked) also completed the analysis in less than 3 h. The maximum memory usage of BWR-finder was 6.53 GB for the TRs masked condition and 6.86 GB for the TRs not masked condition. These values were substantially lower than those of HiTE (92.00 GB) and REPrise (35.76 GB), although RepeatModeler2 required the least memory (1.61 GB). However, the low memory usage of RepeatModeler2 reflects its genome-sampling strategy and is therefore not directly comparable to whole-genome analysis.

**Fig. 4.**
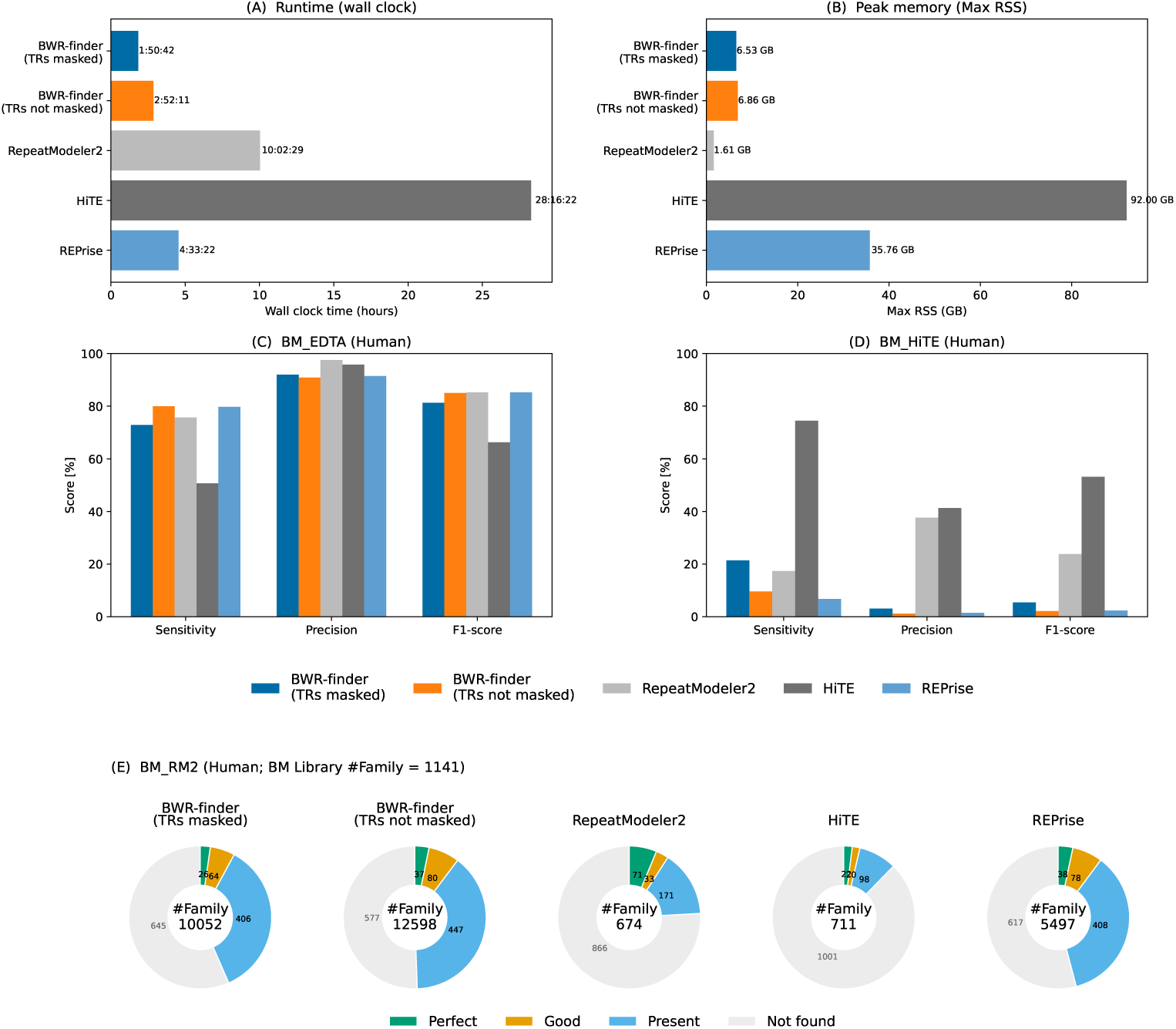
Benchmark results of BWR-finder on the human genome. (A) Runtime. (B) Memory usage. (C) BM EDTA. (D) BM HiTE. (E) BM RM2. The numbers inside the donut holes indicate the number of consensus sequences in each Test Library.

In BM EDTA, BWR-finder (TRs not masked) achieved the highest sensitivity, at 79.93%, followed by REPrise (79.76%), RepeatModeler2 (75.72%), and BWR-finder (TRs masked) (72.89%). HiTE showed lower sensitivity, at 50.65%. Precision was highest for RepeatModeler2, at 97.51%. The other methods also maintained high precision values above 90%, indicating that all methods produced relatively few false-positive repeat regions in this benchmark. The F1-scores of BWR-finder, RepeatModeler2, and REPrise were nearly equivalent, at approximately 85%. These results indicate that BWR-finder detected repeat regions with the highest sensitivity while maintaining false positives at a level comparable to existing methods.

In BM HiTE, HiTE achieved the highest sensitivity, at 74.45%, followed by BWR-finder (TRs masked) (21.38%), RepeatModeler2 (17.37%), BWR-finder (TRs not masked) (9.60%), and REPrise (6.77%). In contrast, the number of FP bases increased markedly for BWR-finder and REPrise (Supplementary Table S6), resulting in very low precision values below 3%. RepeatModeler2, which uses genome sampling, showed a smaller decrease in precision (38%). These results suggest that, as genome size increases, BWR-finder tends to produce a larger number of fragmented repeats.

In the BM RM2 evaluation, RepeatModeler2 produced the largest number of Perfect models (71), followed by REPrise (38), BWR-finder (TRs not masked) (37), BWR-finder(TRs masked)(26) and HiTE (22). In contrast, the cumulative number of sequences classified as Present or better was highest for BWR-finder (TRs not masked) (564), followed by REPrise (524), BWR-finder (TRs masked) (496), RepeatModeler2 (275), and HiTE (140). In particular, the difference between BWR-finder and RepeatModeler2 was larger than that observed for the rice genome (Fig. 4(E)), suggesting that using the whole genome as input improved the recovery of benchmark repeat library sequences. However, as observed for the rice genome, BWR-finder tended to produce an excessive number of repeat consensus sequences: 10,052 sequences in the TRs masked condition and 12,598 sequences in the TRs not masked condition, compared with 1,141 sequences in the benchmark repeat library.

### 2.6 Evaluation on genomes larger than 10 Gb

We evaluated the scalability and detection performance of BWR-finder on large eukaryotic genomes ranging from 14.6 Gb to 87.2 Gb (Fig. 5(A,B)). The total runtime of BWR-finder increased approximately linearly with genome size (Fig. 5(A)), showing a strong correlation ((*R*^2^ = 0.990)). Notably, even for the 87.2-Gb genome, the runtime did not show an exponential increase, indicating stable computational scaling. Similarly, the maximum memory usage showed a strong linear relationship with genome size (Fig. 5(B); (*R*^2^ = 0.978)), indicating predictable memory scaling. These results show that BWR-finder maintains stable computational characteristics even for extremely large genomes.

**Fig. 5.**
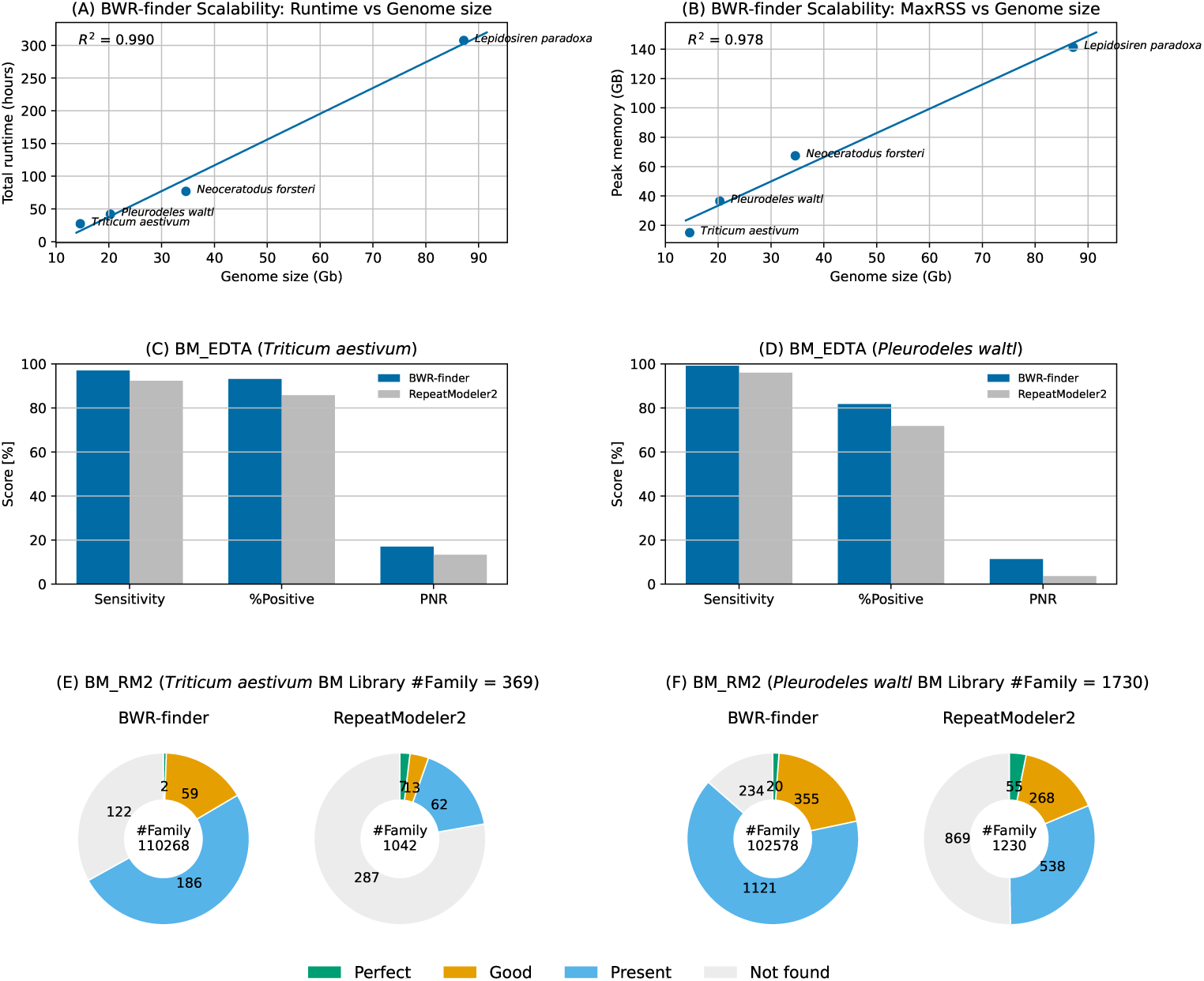
Results for large genomes. (A) Runtime of BWR-finder as a function of genome size. (B) Memory usage of BWR-finder as a function of genome size. (C) BM EDTA results for BWR-finder and RepeatModeler2 on the wheat genome (15 Gb). %Positive indicates the proportion of the whole genome annotated as repeats, and Potential Novel Repeat (PNR) indicates the proportion of the whole genome classified as false-positive regions. (D) BM EDTA results for the salamander genome (20.3 Gb), shown as in (C). (E) BM RM2 results for BWR-finder and RepeatModeler2 on the wheat genome. (F) BM RM2 results for BWR-finder and RepeatModeler2 on the salamander genome.

We next compared detection performance using the BM EDTA metrics, including sensitivity, %Positive, and Potential Novel Repeat (PNR). In wheat (*Triticum aestivum*), BWR-finder showed high sensitivity and recovered a large proportion of the existing repeat annotation (Fig. 5(C)). %Positive and PNR were 93.1% and 17.0%, respectively, indicating that BWR-finder predicted additional repeat regions beyond the existing repeat annotation. Because wheat has a hexaploid genome, some non-TE-derived duplicated regions may have been detected as repeats. Nevertheless, these results indicate that BWR-finder detected repeat regions that were absent from the existing repeat annotation. BWR-finder also maintained high sensitivity in the salamander genome (*Pleurodeles waltl* ) (Fig. 5(D)). Compared with RepeatModeler2,

BWR-finder showed %Positive and PNR values that were approximately 10 percentage points higher. This result suggests an advantage of using the whole genome as input, because this strategy avoids missing repeats located outside sampled genomic regions.

We further compared BWR-finder and RepeatModeler2 using BM RM2 to evaluate repeat library recovery (Fig. 5(E,F)). In both wheat ((n = 369)) and salamander ((n = 1730)), BWR-finder recovered more sequences in the Good and Present categories than RepeatModeler2. These results suggest that BWR-finder improves repeat library recovery in large and highly repetitive genomes. However, RepeatModeler2 recovered more Perfect models, and BWR-finder produced a very large number of detected consensus sequences, exceeding 100,000 sequences. Consistent with the results for the rice and human genomes, these observations indicate that BWR-finder still has a limitation in that its output contains fragmented repeat consensus sequences.

### 2.7 Exploration of previously uncharacterized repeats in *Pleurodeles waltl*

The comparison between the existing repeat annotation and the BWR-finder-based annotation for the salamander (*Pleurodeles waltl* ) genome is shown in Fig. 6(A) and Supplementary Table S4. In the existing repeat annotation, 14.74 Gb of the *Pleurodeles waltl* genome, corresponding to 72.6% of the genome, was annotated as repeats. In contrast, the BWR-finder-based prediction annotated 17.12 Gb, corresponding to 84.4% of the genome, as repeats. Thus, BWR-finder detected an additional 2.39 Gb of repeat regions compared with the existing repeat annotation, corresponding to 11.8% of the genome.

**Fig. 6.**
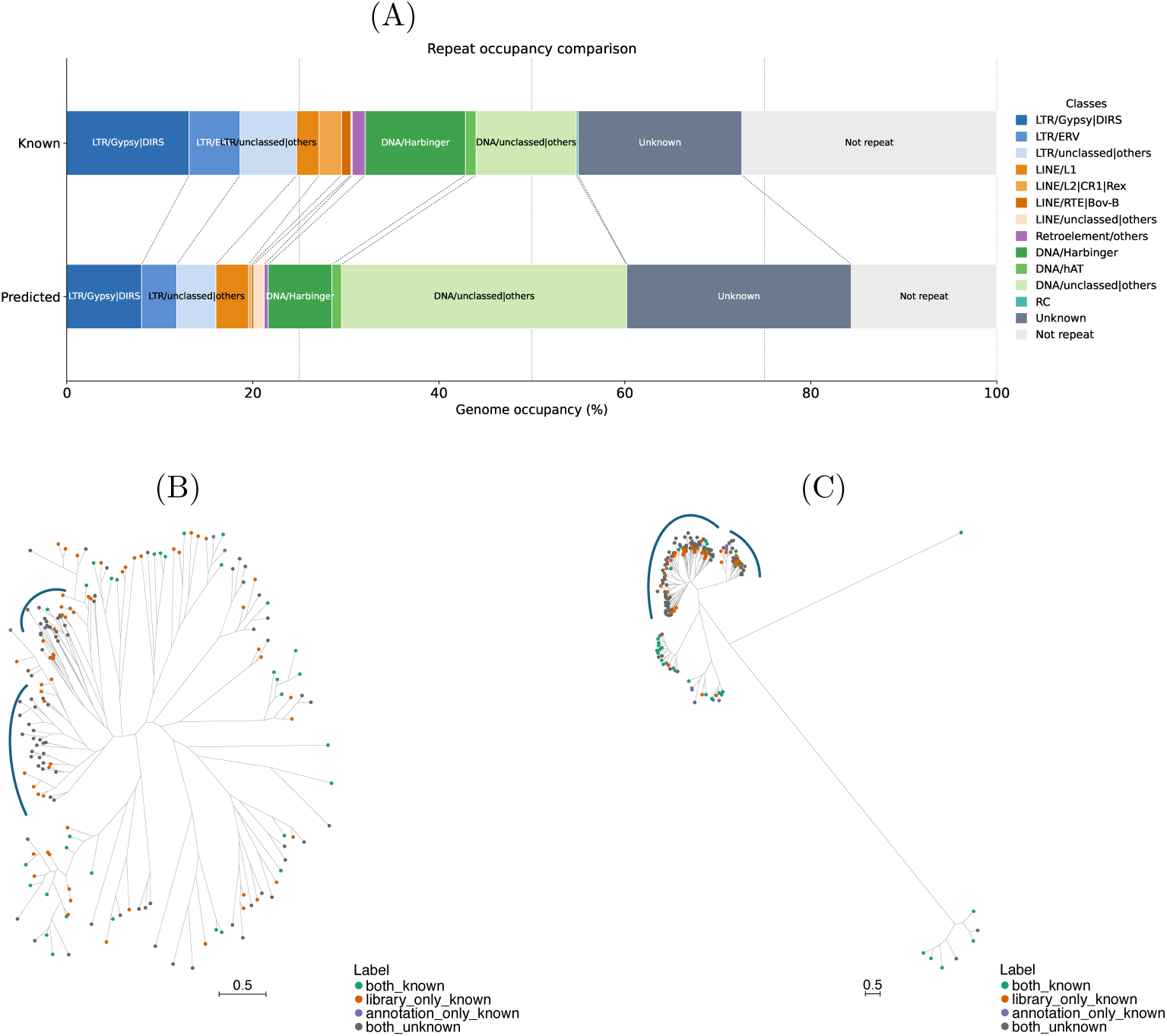
Detailed repeat analysis in *Pleurodeles waltl*. (A) Comparison of RepeatMasker coverage after polishing and classification. Terrier was also applied to the exisiting repeat library, and repeat classes were adjusted according to their proportions in the RepeatMasker.tbl output (Supplementary Table S4). (B) Phylogenetic tree based on nucleotide sequences corresponding to Pol regions extracted from LINE/L1 repeat consensus sequences generated by the BWR-finder-based pipeline, including BWR-finder. Sequences were extracted when at least 80% of their length aligned to LINE/L1-derived Pol protein sequences. Navy markers indicate clades that could not be assigned to known annotated repeats. (C) Phylogenetic tree of nucleotide regions corresponding to Pol proteins from repeat consensus sequences classified as LTR/Gypsy, shown as in (B).

At the repeat-class level, the BWR-finder-based annotation showed lower coverage for several classes, including LTR/Gypsy—DIRS, which decreased from 2.67 Gb (13.1%) to 1.63 Gb (8.04%), DNA/Harbinger, which decreased from 2.19 Gb (10.8%) to 1.39 Gb (6.84%), and LINE/L2—CR1—Rex, which decreased from 486.5 Mb (2.40%) to 62.4 Mb (0.31%). In contrast, greater coverage was observed for LINE/L1, LINE/unclassed—others, DNA/unclassed—others, and Unknown repeats.

Among these classes, the increases were particularly pronounced for DNA/unclassed—others and Unknown repeats. For DNA repeats as a whole, the existing repeat annotation covered 4.61 Gb (22.7%) of the genome, whereas the BWR-finder-based annotation covered 7.82 Gb (38.5%). However, most of this increase was attributable to regions classified as DNA/unclassed—others rather than to regions assigned to specific DNA subclasses. DNA/unclassed—others increased from 2.19 Gb (10.8%) in the existing repeat annotation to 6.22 Gb (30.7%) in the Analysis Annotation. Unknown repeats also increased from 3.57 Gb (17.6%) in the existing snnotation to 4.91 Gb (24.2%) in the Analysis Annotation. Therefore, many repeat regions additionally detected by BWR-finder were observed as unclassed or Unknown repeats rather than as regions clearly assigned to known subclasses.

Fig. 6(B) and (C) show phylogenetic trees based on nucleotide regions corresponding to Pol proteins extracted from BWR-finder-derived LINE/L1 and LTR/Gypsy repeat consensus sequences, respectively. Here, library-known and annotation-known indicate consensus sequences showing at least 80% coverage against the existing repeat library and existing repeat annotation, respectively (Supplementary Figure S3). In both repeat classes, BWR-finder-derived consensus sequences that met neither criterion formed large clades in the phylogenetic trees. These clades suggest that BWR-finder detected groups of related repeat consensus sequences that were not represented in the existing repeat library or annotation.

## 3 Discussion

In this study, we proposed a method for low-memory and practical *de novo* repeat library construction for large genomes by using a BWT-based seed-and-extend approach. In conventional methods based on suffix arrays or *k*-mer occurrence tables, the increase in memory usage with genome size has been a major limitation for application to large genomes. In contrast, the proposed method manages repeat occurrences on the BWT rather than as coordinates on the original sequence and performs extension using LF/FL mapping. This design enables repeat sequences to be detected while keeping memory usage low. Our results show that a BWT-based seed-and-extend method is a practical approach for *de novo* repeat detection in large and repeat-rich genomes exceeding 10 Gb.

Although BWR-finder outperformed existing methods in terms of speed and memory usage, fragmentation of the generated repeat library remains a limitation. When the detected repeat library is used not only for genome masking but also for repeat classification or evolutionary analysis, fragmentation of candidate sequences becomes an important issue. One possible cause of fragmentation is that the proposed method does not use the frequency-based seed-processing order adopted by REPrise and RepeatScout. Incorporating such an extension order would require additional seed information in a priority queue and would also make extension operations less independent during parallel computation. Therefore, the fragmentation of repeat consensus sequences in the library generated by BWR-finder reflects a technical trade-off between computational efficiency, including runtime and memory usage, and repeat-library quality.

However, fragmentation is not a limitation specific to the proposed method. In seed-and-extend-based *de novo* repeat detection, it is inherently difficult to determine complete repeat boundaries by extension and multiple sequence alignment when copies are partially deleted, highly diverged, or nested within other repeats [28]. Indeed, our analyses of the rice and human genomes showed that all seed-and-extend tools produced fragmented repeat sequences to varying degrees. One practical solution to this problem is to combine sensitive repeat sequence detection with subsequent library polishing. In the salamander analysis, post-processing reduced the number of candidate sequences by approximately half, suggesting that redundant or fragmented candidate sequences can be partially integrated after detection. From this perspective, the proposed method is useful as an upstream step that efficiently generates comprehensive repeat candidates from large genomes.

Another practical consideration is whether TRs should be masked before applying the proposed method. In the current extension procedure, genomic distances among seed occurrences are not explicitly used, making it difficult to distinguish TRs from interspersed repeats during extension. Pre-masking TRs can reduce computational cost and false-positive repeats. Therefore, in the analysis pipeline used in Section 2.7, we used TR-masked genomes as input. However, in the rice and human genomes, pre-masking reduced sensitivity by approximately 5 %. Thus, the masking strategy should be selected according to the purpose of the analysis. For conservative repeat annotation, pre-masking is effective for reducing false positives. In contrast, when comprehensive detection of interspersed repeats is the primary goal, an unmasked genome can be used as input, followed by downstream polishing and classification to remove sequences derived from TRs.

Our experiments also showed that the bottleneck in repeat analysis of large genomes is not limited to de novo library construction. For the *Pleurodeles waltl* genome, RepeatMasker required 17days, and TEstrainer could not directly process the entire library because of memory limitations; therefore, the input had to be split into batches of 1,000 sequences (Supplementary Table S7). Thus, future large-genome repeat analysis will require not only faster and more memory-efficient *de novo* detection but also the development of downstream methods that can be applied to large-scale data.

For downstream method development, our algorithm suggests that BWT-based representations may be useful beyond the repeat detection step. In repeat-rich genomes, characters with similar sequence contexts tend to cluster on the BWT, and BWT-based compressed indexes can store and search such sequences efficiently. Therefore, introducing BWT-based analytical strategies into downstream processes may help reduce the computational requirements of the entire repeat analysis pipeline for large genomes.

## 4 Conclusion

In this study, we aimed to detect interspersed repeats in large genomes with lower memory usage and practical runtime. We developed a seed-and-extend-based method for *de novo* interspersed repeat detection that manages each seed occurrence and its surrounding regions using the BWT/FM-index. The proposed method substantially reduced computational resource requirements while maintaining detection performance. These results indicate that BWT-based repeat detection can provide a practical framework for analyzing large and repeat-rich genomes.

## 5 Methods

### 5.1 Overview of the method

We propose a method for detecting interspersed repeats in which a genome sequence is represented using the BWT and the FM–index. Seed search, seed extension, and masking are all performed on the BWT sequence (Fig. 1(B)). In conventional seed-and-extend methods, each seed occurrence is retained as a coordinate on the original sequence, and the neighboring bases around each seed occurrence must be accessed during the extension step. In contrast, our method manages each seed occurrence on the BWT sequence, and the adjacent bases are retrieved by LF/FL mapping on the BWT sequence during the extension step. This design allows not only seed search but also extension to be completed on the BWT sequence.

The overall procedure is shown in Fig. 1(B) and Supplementary Algorithm S5. First, frequent *k*-mer positions are enumerated as seeds on the FM-index. Next, each set of seed occurrences is extended in the left and right directions, and consensus sequences are constructed iteratively based on alignment scores. Finally, positions on the BWT corresponding to detected repeat regions are masked to prevent the same regions from being repeatedly selected in subsequent seed searches. In this method, this series of operations is handled uniformly as interval and row operations on the BWT.

### 5.2 Representation of sequence occurrences on the BWT

Let *T* be a genome sequence of length *n* with a unique terminal symbol $ appended, and let *SA* be its suffix array. The BWT is defined as the array *L*, which consists of the characters immediately preceding the suffixes ordered by the suffix array:

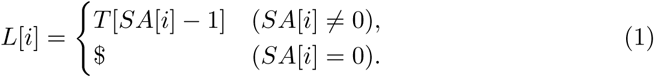

In this method, all occurrences of a string pattern *P* are represented as the half-open interval [*l, r*) on the suffix array that corresponds to the set of suffixes having *P* as a prefix. Thus, the occurrence set of a seed can be managed as an interval on the BWT sequence without explicitly retaining individual genomic coordinates.

The first column *F* in the BWT sequence corresponds to the stable sorting of the characters in *L*. Using this property, LF mapping can be defined as a mapping from a position *i* on the BWT to the row corresponding to the position one character to the left on the original sequence:

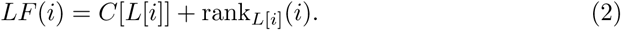

Here, *C*[*c*] denotes the total number of characters lexicographically smaller than character *c*, and rank*_c_*(*i*) denotes the number of occurrences of character *c* in the array *L*[0*..i*). Similarly, FL mapping is defined as the mapping from a position in the first column *F* to the corresponding position on the BWT sequence:

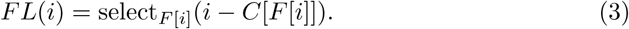

Here, select*_c_*(*k*) returns the position of the *k*-th occurrence of character *c*. In this method, movement in the left and right directions on the original sequence is realized on the BWT sequence by updating BWT rows using these mappings. Therefore, in addition to seed search, access to adjacent bases required for extension can also be performed using only operations on the BWT sequences.

A schematic illustration of the BWT and FM-index, together with examples of LF/FL mapping, is shown in Supplementary Figure S1.

### 5.3 Seed search on the BWT

Seeds are searched using an FM-index backward search with frequency-based pruning. In the FM-index, the search state is maintained as a suffix array interval [*l, r*). When a character *a* ∈ *A, C, G, T* is prepended to the current string, the corresponding interval is updated as

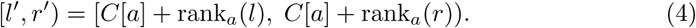

By repeating this interval update, the occurrence set of a pattern can be tracked on the BWT sequence without explicitly retaining the suffix array or individual occurrence positions. In this method, to enumerate frequent *k*-mers as seeds, the search starts from the root node (*ε,* [0*, n*)), which corresponds to the empty string, and proceeds in a depth-first manner by extending the string one character at a time. When a node reaches length *k*, the corresponding *k*-mer is adopted as a seed candidate if its effective frequency is greater than or equal to the threshold *f*_thr_. The detailed procedure is shown in Supplementary Algorithm S1.

To improve search efficiency, for each interval [*l, r*), the effective frequency EffectiveFreq([*l, r*), Mask) is computed by subtracting the number of masked occurrences from the interval width *r*−*l*. Branches for which this value is below the threshold are terminated because they are not expected to generate seeds with sufficient frequency. This pruning suppresses searches for seeds that strongly correspond to already detected regions and for seed candidates with low occurrence counts.

### 5.4 Extension on the BWT sequence

#### 5.4.1 Consensus-based extension

The extension procedure in this method follows the consensus extension framework introduced in RepeatScout[12]. Specifically, for a set of sequences *S*_1_*, . . . , S_M_* obtained from the regions near the endpoints of the seed occurrences, the procedure is formulated as a problem of sequentially extending a partial consensus sequence *Q t* while maximizing a multiple alignment score *A*(*Q*; *S*_1_*, . . . , S_M_* ).

At the *t*-th extension step, candidates *Q_t_*· *N* , obtained by appending one base *N* ∈ *A, C, G, T* to the end of *Q_t_*, are evaluated, and the next consensus sequence is determined as

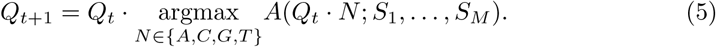

Here, *A*(·) is defined based on local alignment scores against each occurrence sequence. We followed the scoring scheme of REPrise[13] and reimplemented the extension process so that it is compatible with occurrence management on the BWT sequence.

The stopping criterion for extension follows the best-scoring prefix, and terminates extension when the score does not improve for a predefined number of steps. This suppresses unstable terminal extension while estimating representative sequences for repeats. The details of right extension are shown in Supplementary Algorithm S2. Left extension is performed symmetrically. See Supplementary Methods and Supplementary Table S3 for details.

#### 5.4.2 Retrieving adjacent bases via LF/FL mapping

In this method, each occurrence of a seed is retained as a BWT row, rather than as a coordinate on the original sequence. Therefore, to access the adjacent base for each occurrence at each extension step, the corresponding row on the BWT must be updated instead of simply updating a position on the original sequence. For this purpose, our method uses LF/FL mapping to move to the BWT row corresponding to an adjacent position on the original sequence.

For right extension, FL mapping updates the BWT row for each occurrence, corresponding to a one-base movement to the right on the original sequence. The base is then retrieved from the destination row and used as the next base for each occurrence. By repeating this operation for all occurrences, the set of bases required at each extension step can be updated on the BWT sequence. As shown in Supplementary Algorithm S3, this method retains the BWT row and the corresponding window for each occurrence and updates them synchronously during extension. Left extension can be performed symmetrically by applying LF mapping instead of FL mapping. Thus, extension of consensus sequences in both directions is performed consistently as mapping operations on the BWT sequence.

### 5.5 Masking on the BWT

Masking is performed to prevent repeat regions that have already been detected from being selected again in subsequent seed searches. Because this method manages seed occurrences and extension on the BWT, masking must also be handled at the level of the BWT.

Specifically, for each character *c* in the character set {*A, C, G, T* }, a bit vector

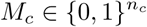

is prepared for all occurrences of that character. Here,

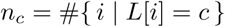

is the total number of occurrences of character *c* on the BWT sequence. When character *c* = *L*[*i*] appears at position *i* on the BWT sequence, the occurrence is recorded as masked by setting

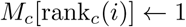

using its occurrence rank rank *c*(*i*). Detected repeat regions are given as a set of pairs of points (*from, to*) on the BWT sequence. In this method, the corresponding BWT rows are sequentially scanned by following LF mapping between these points, and the bit corresponding to each position is updated to 1. The detailed procedure is shown in Supplementary Algorithm S4.

After mask updates, the rank support of the bit vector is reconstructed for each *M_c_*. This allows the number of masked occurrences included in any interval [*l, r*) to be obtained in constant time. This value is used to compute EffectiveFreq during seed search and is reflected in search pruning as described above.

### 5.6 Implementation details

In this implementation, we used the wavelet tree library in sdsl-lite [23] to efficiently support rank and select operations on the BWT sequence. Because DNA sequences have a small number of characters(*σ* = 4), those operations on the wavelet tree can be performed in effectively constant time.

We also used a run-length compressed BWT to reduce memory usage. However, the algorithmic description in this section is based on the standard BWT representation, and the theoretical discussion is presented on that basis. The BWT sequence was constructed using the linear-time and linear-space algorithm implemented in grlBWT [22].

### 5.7 Analysis of repeats detected in the *Pleurodeles waltl* genome

Based on the output of BWR-finder, we constructed a more refined repeat library and repeat annotation for *Pleurodeles waltl* (Fig.1(A)). First, the repeat sequences detected by BWR-finder were polished and classified. For polishing, we ran one round of TEstrainer, which is used in EarlGrey [25]. After clustering the polished sequences using CD-HIT, we classified them using Terrier [26] and constructed the Analysis Library.

Next, we ran RepeatMasker on the *Pleurodeles waltl* genome using the Analysis Library and generated the Analysis Annotation. To compare this annotation with the BWR-finder-derived annotation, we also classified Unclassified repeats in the existing *Pleurodeles waltl* annotation using Terrier, following Brown et al. [36], and incorporated the classification results into the existing repeat annotation (Supplementary Figure S3(A)).

We then compared the existing repeat library and annotation with the BWR-finder-derived Analysis Library and Analysis Annotation in *Pleurodeles waltl*. First, for both the existing repeat annotation and the Analysis Annotation, we calculated the genomic occupancy of each repeat class. Next, consensus sequences in the Analysis Library were classified according to two criteria: (A) whether the coverage of the Analysis Library against the existing repeat annotation was at least 80%, in a BM EDTA-like manner, and (B) whether the coverage in the RepeatMasker alignment between the existing repeat library and the Analysis Library was at least 80%, in a BM RM2-like manner. Based on these two criteria, each sequence in the Analysis Library was assigned one of four labels: annotation known, library-known, both known, or Unknown (Supplementary Figure S3(B)).

For the labeled sequences in the Analysis Library, we performed BLASTX [29] searches for each repeat class using the RepeatPeps database included with Repeat-Masker 4.2.1. Among these sequences, Among these sequences, we extracted nucleotide regions that aligned to Pol proteins over at least 80% of their length. Finally, for each repeat class, we used the labeled Pol-region sequence derived from consensus sequences as input and constructed phylogenetic trees using MAFFT [38], trimAl [39], and IQ-TREE [40] (Supplementary Figure S3(C)).

## Supplementary information

## Acknowledgements

We thank the members of Hamada Laboratory, especially Kento Kubo and Masato Kosuge for their valuable comments. A.T. is grateful to Sigehiro Kuraku (National Institute of Genetics) and Yohei Ogawa (National Institute of Genetics) for insightful discussions. Computations were partially performed on the NIG supercomputer at ROIS National Institute of Genetics.

## Declarations

### Ethics approval and consent to participate

Not applicable. This study used publicly available genome sequence resources and did not involve new human participants, human samples, or animal experiments.

### Consent for publication

All authors consent to the publication of this manuscript.

### Data and Materials availability

Repeat annotations predicted for the *Pleurodeles waltl* genome using BWR-finder are available at https://waseda.box.com/v/BWR-finder-data.

### Code availability

The source code of BWR-finder and scripts for evaluation are freely available at https://github.com/hmdlab/BWR-finder.

### Author contribution

A.T. conceived and proposed the study, implemented the software, developed the evaluation pipeline, performed the experiments and evaluations, and analyzed the results. T.F. and H.M. supervised the study. A.T. prepared the original draft of the manuscript, and T.F. and H.M. reviewed and revised it. All authors contributed to the study and approved the final manuscript.

