## Supplementary Materials for "BWR-finder: BWT-based *de novo* interspersed repeat detection for gigantic genomes"

### S1 Supplementary Methods

#### S1.1 Alignment scoring and termination criterion used in BWR-finder

BWR-finder adopts the alignment-scoring formulation used in REPrise for consensus extension and masking, while replacing the underlying sequence search and masking operations with BWT/FM-index-based procedures. This section summarizes the scoring scheme used in BWR-finder.

During extension, a seed sequence is extended one nucleotide at a time in both directions. For the rightward extension, let $Q_t$ denote the partial consensus sequence after $t$ extension steps, and let $S_1, \ldots, S_M$ denote the genomic sequences adjacent to the seed occurrences. At each step, BWR-finder selects the nucleotide $N \in A, C, G, T$ that maximizes the alignment score

$$A(Q_t \cdot N; S_1, \ldots, S_M),$$

and sets

$$Q_{t+1} = Q_t \cdot N.$$

The extension is terminated when the best-scoring position $t_{\max}$ has not been updated for $w$ consecutive extension steps. The value of $t_{\max}$ is updated to $t$ when

$$A(Q_t; S_1, \ldots, S_M) \geq A(Q_{t_{\max}}; S_1, \ldots, S_M) + r(t - t_{\max}),$$

where $r$ is the penalty for extending genomic regions as repeat instances. Following REPrise and RepeatScout, $r$ was set to 3. In this study, $w$ was set to 100. After termination, $Q_{t_{\max}}$ is concatenated to the corresponding side of the seed sequence. The same procedure is applied independently in the leftward direction, yielding the final consensus sequence $Q$.

The alignment score is computed using banded dynamic programming with affine gap penalties. For two sequences $X$ and $Y$, three DP tables, $M$, $I_X$, and $I_Y$, are updated within the band $i - b \leq j \leq i + b$ as follows:

$$M(i,j) = \max \begin{cases} M(i-1, j-1) + \mu_{ij}, \\ I_X(i-1, j-1) + \mu_{ij}, \\ I_Y(i-1, j-1) + \mu_{ij}, \end{cases}$$

$$I_X(i,j) = \max \begin{cases} M(i-1, j) + o, \\ I_X(i-1, j) + e, \end{cases} \qquad I_Y(i,j) = \max \begin{cases} M(i, j-1) + o, \\ I_Y(i, j-1) + e, \end{cases}$$

where $1 \leq i \leq |X|$, $1 \leq j \leq |Y|$, $b$ is the band width, $o$ is the gap-opening score, and $e$ is the gap-extension score. The substitution score $\mu_{ij}$ is $+1$ for a match and $-1$ for a mismatch between the $i$-th character of $X$ and the $j$-th character of $Y$. The alignment score at position $(i, j)$ is defined as

$$a(i,j) = \max\{M(i,j), I_X(i,j), I_Y(i,j)\}.$$

In the masking step, BWR-finder determines the genomic region corresponding to the aligned portion of the final consensus sequence $Q$. For rightward masking, let $a_{Q,S_m}(i,j)$ be the alignment score between the first $i$ characters of $Q$ and the first $j$ characters of $S_m$. If $(i', j')$ gives the maximum value of $a_{Q,S_m}(i,j)$, the region corresponding to the first $j'$ characters of $S_m$, together with the seed region, is masked. The same procedure is applied in the leftward direction. In BWR-finder, these masking operations are implemented on the BWT representation of the genome using nucleotide-specific bit vectors.

#### S1.2 Datasets and reference annotations

The genome assemblies, repeat libraries, and reference annotations used in this study are summarized in Table S1. For genomes used only in scalability experiments, reference libraries and annotations were not used.

#### S1.3 Software versions and container images

The software versions and container images used in this study are listed in Table S2. For tools installed from source code, commit hashes are reported where available.

#### S1.4 Parameter settings

The major parameters used for BWR-finder and post-processing are summarized in Table S3. Unless otherwise specified, experiments were run using 32 CPU threads.

Table S1: Genome resources used in this study.

| Resource | Genome | Repeat library | Annotation |
|---|---|---|---|
| Rice | MSU7<br>from EDTA database[6] | rice7.0.0.liban<br>from EDTA database | Rice_MSU7.fasta.std7.0.0.out<br>from EDTA database |
| Human | chm13v2.0.fa<br>from T2T consortium[5] | Repbase 26.02 [1]<br>human_rep + human_sub | chm13v2.0.fa repeat BED file[3].<br>thickStart and thickEnd columns |
| Wheat | GCF_018294505.1_<br>IWGSC_CS_RefSeq_v2.1<br>genomic.fna[9]<br>from UCSC genome browser | trep-db_nr_Rel-25<br>Triticum_aestivum.fasta<br>from TREP[8] | GCF_018294505.1_<br>IWGSC_CS_RefSeq_v2.1<br>repeatMasker.out<br>from UCSC genome browser |
| Salamander | GCA_026652325.1.fa.gz[2] | GCA_026652325.1<br>rmsk.customLib.fa.gz<br>from UCSC genome browser | GCA_026652325.1<br>repeatModeler.out.gz<br>from UCSC genome browser |
| N. forsteri<br>Lungfish<br>(34.6 Gb) | GCA_016271365.2_neoFor_v3.1<br>genomic.fna[4] | – | – |
| L. paradoxa<br>Lungfish<br>(87.2 Gb) | GCA_040581445.1_<br>ASM4058144v1 genomic.fna[7] | – | – |

Table S2: Software/library versions and container images used in this study.

| Software/library | Version / commit | Purpose |
|---|---|---|
| BWR-finder | 1.0.0 | Proposed de novo repeat detection |
| grlBWT | 1.0.1 alpha | C++ Library for BWT construction |
| sdsl-lite | 2.1.0 | C++ Library for Succinct data structures |
| REPrise | 1.0.1 | Comparison |
| RepeatModeler2 | 2.0.7 | Comparison |
| HiTE | 3.3.3 | Comparison |
| RepeatMasker | 4.2.1 | Genome masking |
| CD-HIT | 4.8.1 | Clustering repeat libraries |
| TEstrainer | commit `6a225c5` | Polish repeat libraries |
| earlgrey | 6.3.6 | Container for TEstrainer |
| tantan | 26 | Tandem repeat masking |
| Terrier | 0.3.4 | Repeat classification |
| bedtools | 2.31.1 | Coverage calculation |
| BLAST | 2.17.0+ | Get Pol gene regions from consensus repeats |
| mafft | 7.526 | Multiple sequence alignment of consensus repeats |
| trimAl | 1.5.rev1 | Trim gaps of MSA |
| iqtree | 3.1.1 | Draw phylogenetic tree |

Table S3: Major parameter settings used in this study. $N$ is the input sequence length.

| Step | Parameter | Value | Notes |
|---|---|---|---|
| Seed search | Seed length $k$ | $\log_4 N + 1$ | Rice:15, Human:17, Wheat:18, salamander:18, lungfish: 19 |
| Seed search | Frequency threshold $f_{\min}$ | 10 | Minimum effective occurrence count |
| Extension | MAXEXTEND | 10000 | Maximum extension length |
| Extension | WHEN_TO_STOP $w$ | 100 | Stopping criterion |
| Extension | MINIMPROVEMENT $r$ | 3 | Minimum score improvement |

Table S4: Harmonization rules for repeat class labels used to compare the BWR-finder-based analysis annotation and the Terrier-reclassified known annotation in *Pleurodeles waltl*. The analysis annotation was generated by aligning the analysis library to the genome using RepeatMasker. For the known annotation, repeat entries annotated as Unknown or Unclassified were reclassified using Terrier before class harmonization. RepeatMasker `.tbl` labels correspond to rows in the summary table used for occupancy calculations, whereas RepeatMasker `.out` labels correspond to raw class labels in the annotation file. Asterisks indicate prefix matching.

| Harmonized class used in this study | RepeatMasker `.tbl` label | RepeatMasker `.out` class label |
|---|---|---|
| SINE | `SINEs` | `SINE`, `SINE/*` |
| PLE | `Penelope` | `PLE`, `PLE/*`, `Penelope`, `Penelope/*`, `LINE/Penelope` |
| LINE/L1 | `L1/CIN4` | `LINE/L1`, `LINE/L1-Tx1` |
| LINE/L2\|CR1\|Rex | `L2/CR1/Rex` | `LINE/L2`, `LINE/CR1`, `LINE/Rex-Babar` |
| LINE/R1\|LOA\|Jockey | `R1/LOA/Jockey` | `LINE/I`, `LINE/I-Jockey`, `LINE/R1`, `LINE/LOA` |
| LINE/R2\|R4\|NeSL | `R2/R4/NeSL` | `LINE/R2`, `LINE/R2-NeSL`, `LINE/R2-Hero` |
| LINE/RTE\|Bov-B | `RTE/Bov-B` | `LINE/RTE`, `LINE/RTE-BovB`, `LINE/RTE-X`, `LINE/RTE-RTE` |
| LINE/unclassed\|others | Residual of `LINEs` total after subtracting explicit LINE child rows | `LINE`, other `LINE/*` labels |
| LTR/Copia | `Ty1/Copia` | `LTR/Copia` |
| LTR/Gypsy\|DIRS | `Gypsy/DIRS1` | `LTR/Gypsy`, `LTR/DIRS`, `LTR/Ngaro` |
| LTR/ERV | `Retroviral` | `LTR/ERV`, `LTR/ERV*` |
| LTR/unclassed\|others | `BEL/Pao`; residual of `LTR elements` total after subtracting explicit LTR child rows | `LTR`, other `LTR/*` labels |
| DNA/hAT | `hobo-Activator` | `DNA/hAT`, `DNA/hAT-*` |
| DNA/TcMar | `Tc1-IS630-Pogo` | `DNA/TcMar`, `DNA/TcMar-*` |
| DNA/CMC | `En-Spm` | `DNA/CMC`, `DNA/CMC-*` |
| DNA/MULE | `MULE-MuDR` | `DNA/MULE`, `DNA/MULE-*` |
| DNA/PiggyBac | `PiggyBac` | `DNA/PiggyBac`, `DNA/PiggyBac-*` |
| DNA/Harbinger | `Tourist/Harbinger` | `DNA/PIF-Harbinger`, `DNA/Harbinger` |
| DNA/unclassed\|others | `Other (Mirage, P-element, Transib)`; residual of `DNA transposons` total after subtracting explicit DNA child rows | `DNA`, other `DNA/*` labels |
| RC | `Rolling-circles` | `RC`, `RC/*` |
| Unknown | `Unclassified` | `Unknown` |
| Excluded | `Simple_repeat`, `Low_complexity`, `Satellite`, `rRNA`, `tRNA`, `snRNA`, `scRNA`, `srpRNA`, `ARTEFACT`, blank labels | `Simple_repeat`, `Low_complexity`, `Satellite`, `rRNA`, `tRNA`, `snRNA`, `scRNA`, `srpRNA`, `ARTEFACT`, blank labels |
| Unmapped | Not assigned to a harmonized class | Other labels not matching any rule |

#### S1.5 Computational environment

Large-scale experiments were conducted on a Linux high-performance computing cluster. Runtime was measured as wall-clock time, and memory usage was recorded as the maximum resident set size. Representative Slurm resource settings are shown below.

Listing 1: Representative Slurm resource settings.

```
#SBATCH --ntasks=1
#SBATCH --cpus-per-task=32
#SBATCH --mem-per-cpu=8G
#SBATCH --time=28-00:00:00
```

#### S1.6 Representative command lines

This section provides representative commands used in the experiments.

##### S1.6.1 Running the proposed method

Listing 2: Representative command for the proposed method.

```
./BWR-finder \
  --genome genome.fa \
  --seed-length XX \
  --min-frequency 10 \
  --threads 32 \
  --output BWR-finder_raw_library.fa
```

##### S1.6.2 Clustering with CD-HIT

Listing 3: Representative CD-HIT command.

```
cd-hit-est \
  -i BWR-finder_raw_library.fa \
  -o BWR-finder_cdhit_c0.8.fa \
  -c 0.8 \
  -p 1 \
  -T 32 \
  -M 0
```

##### S1.6.3 Tandem repeat masking with tantan

Listing 4: Representative tantan command.

```
tantan genome.fa > genome.tantan_masked.fa
```

##### S1.6.4 Genome masking with RepeatMasker

Listing 5: Representative RepeatMasker command.

```
RepeatMasker \
  -pa 32 \
  -no_is \
  -norna \
  -nolow \
  -div 40 \
  -cutoff 225 \
```

```
  -lib BWR-finder_cdhit_c0.8.fa \
  genome.fa
```

##### S1.6.5 Library polishing with TEstrainer

Listing 6: Representative TEstrainer command.

```
./TEstrainer \
  -g genome.fa \
  -l BWR-finder_cdhit_c0.8.fa \
  -t 8 \
  -r 1 \
  -C \
  -d testrainer_out
```

For large libraries, the input library was split into chunks of 1000 sequences and processed independently.

Listing 7: Representative command for splitting a large repeat library.

```
python split_fasta.py \
  --input BWR-finder_cdhit_c0.8.fa \
  --records-per-file 1000 \
  --output-dir split_1000
```

##### S1.6.6 Evaluation

Listing 8: Representative evaluation command.

```
python eval_repeat_bulk.py eval_def.txt > evaluation_results.csv
```

##### S1.6.7 BLASTX search for Pol protein regions

Listing 9: Representative BLASTX command for identifying Pol protein regions in class-specific consensus sequences. Class-specific consensus sequences were searched against class-specific primary RepeatPeps protein subsets. BLASTX hits were first collected with E-value <= 1e-3, and the best hit for each query was selected by bitscore. Hits were retained when E-value <= 1e-5, bitscore >= 50, and subject protein coverage >= 0.80.

```
makeblastdb
-in CLASS.primary.peps.fa
-dbtype prot
-out CLASS.primary.peps

blastx
-query CLASS.consensus.fa
-db CLASS.primary.peps
-out CLASS_vs_primary.blastx.tsv
-outfmt "6␣qseqid␣sseqid␣pident␣length␣mismatch␣gapopen␣qstart␣qend␣sstart␣send␣evalue␣bitscore␣qframe␣
     slen"
-evalue 1e-3
-max_target_seqs 10
-num_threads 4

sort -k1,1 -k12,12nr CLASS_vs_primary.blastx.tsv
| awk '!seen[$1]++' \

> CLASS_vs_primary.besthit.tsv
```

```
awk 'BEGIN{OFS="\t"}
$11␣<=␣1e-5␣&&␣$12␣>=␣50␣&&␣$14␣>␣0␣&&␣($4␣/␣$14)␣>=␣0.80␣{
print␣$0,␣($4␣/␣$14)
}' CLASS_vs_primary.besthit.tsv \

> CLASS_vs_primary.besthit.filt.tsv
```

##### S1.6.8 Construction of Pol-region phylogenetic trees

Listing 10: Representative commands for constructing Pol-region phylogenetic trees. Strand-aware Pol-region sequences extracted from consensus sequences were aligned using MAFFT, trimmed using trimAl, and used for phylogenetic tree inference with IQ-TREE 3. IQ-TREE was run with ModelFinder and 1,000 ultrafast bootstrap replicates. Analyses were performed using four threads, and classes with fewer than three sequences after trimming were excluded from tree inference.

```
mafft
--thread 4
--auto
CLASS.hit_regions_strand.fa \

> CLASS.aln.fa

trimal
-in CLASS.aln.fa
-out CLASS.trim.fa
-automated1

iqtree3
-s CLASS.trim.fa
-m MFP
-bb 1000
-nt 4
-redo
--prefix CLASS
```

#### S2 Supplementary Figures

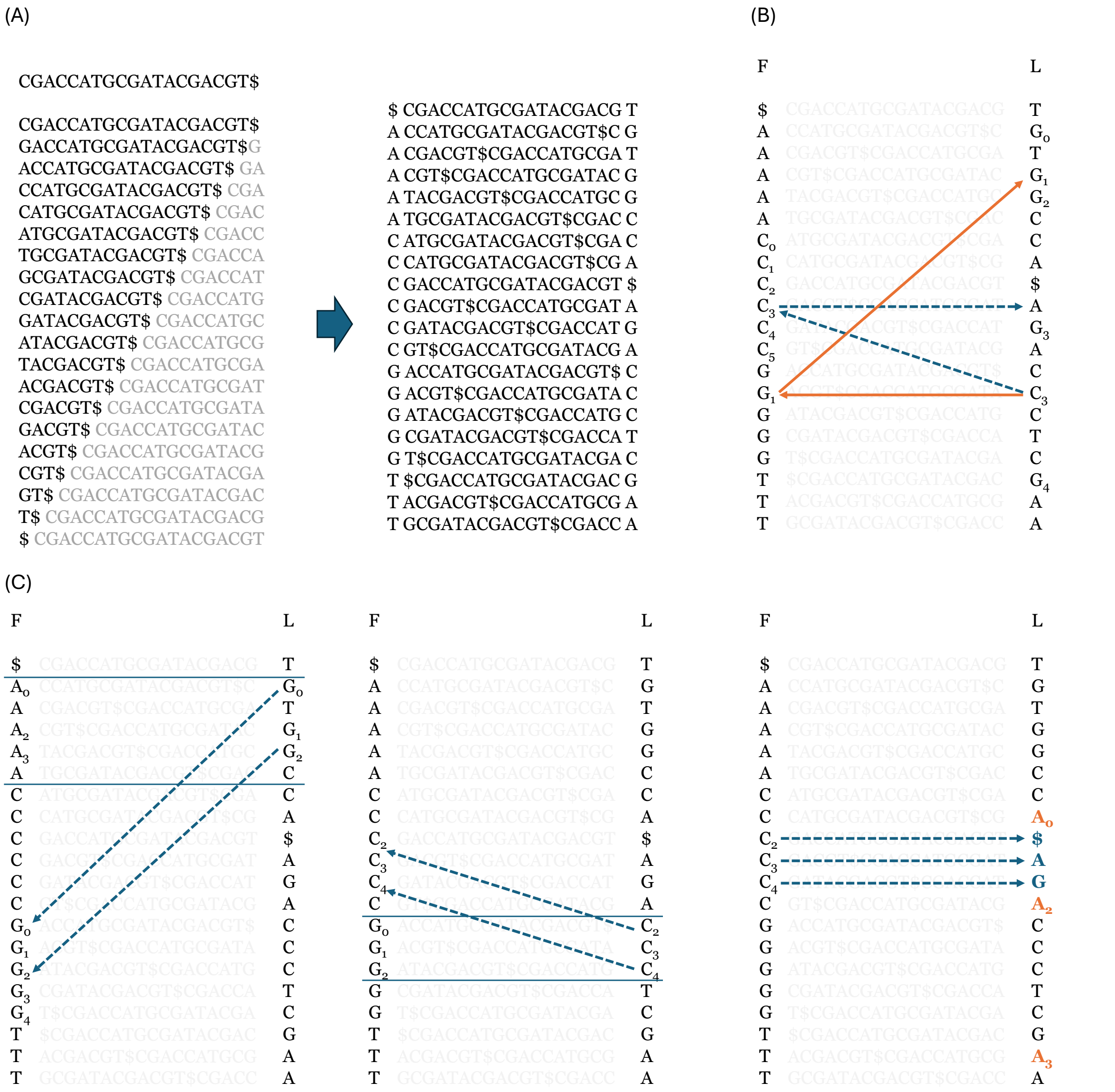

Figure S1: Schematic overview of the BWT and FM-index operations used in BWR-finder. **(A)** Relationship between the suffix array and BWT. **(B)** Example of LF/FL mapping. **(C)** Identification of seed occurrences using FM-index backward search.

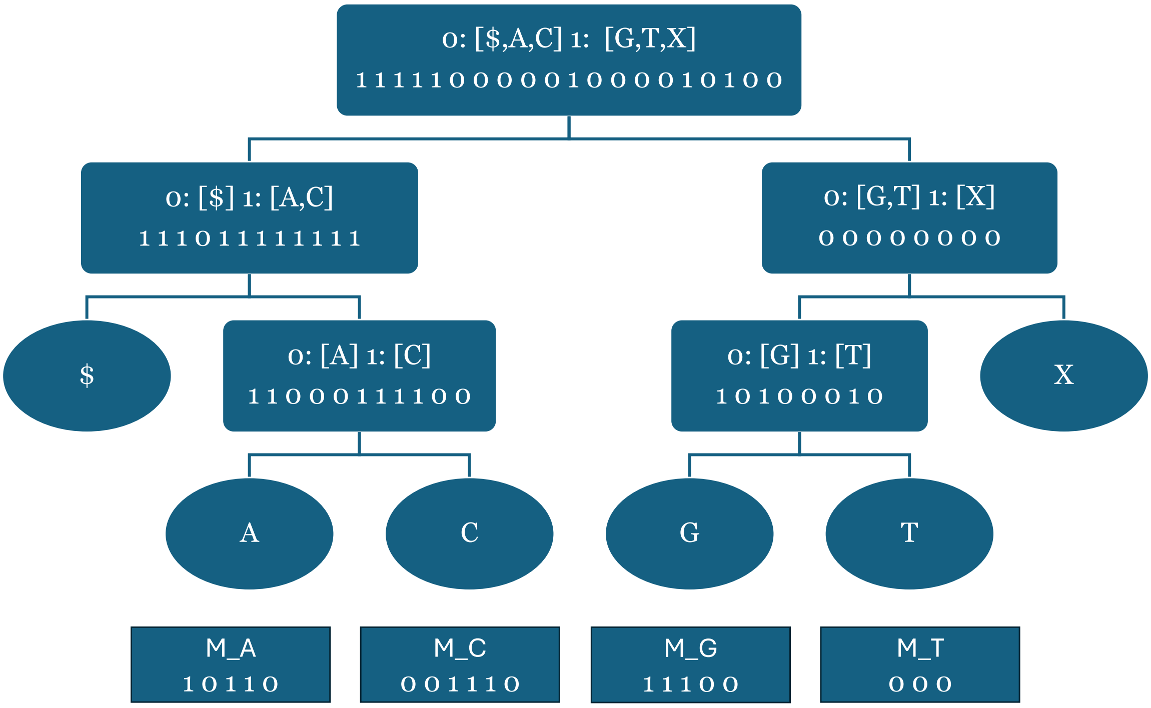


Figure S2: Schematic representation of the wavelet tree and masking bit vector data structures constructed by BWR-finder for the BWT sequence `TGTGGCCA$AGACCCTCGAA`. Each nucleotide character, $A$, $T$, $G$, and $C$, is associated with a masking bit vector whose length equals the number of occurrences of $c$ in the BWT sequence. These bit vectors prevent redundant computation when subsequent seeds match repeat regions that have already been detected. The figure shows the masking state after the seed `CGA`, identified in Fig.S1, has been masked. The symbol \$ denotes the terminal character, whereas $X$ represents regions excluded from repeat detection, such as tandem repeat regions and chromosome boundaries. Neither \$ nor $X$ is used for repeat detection.

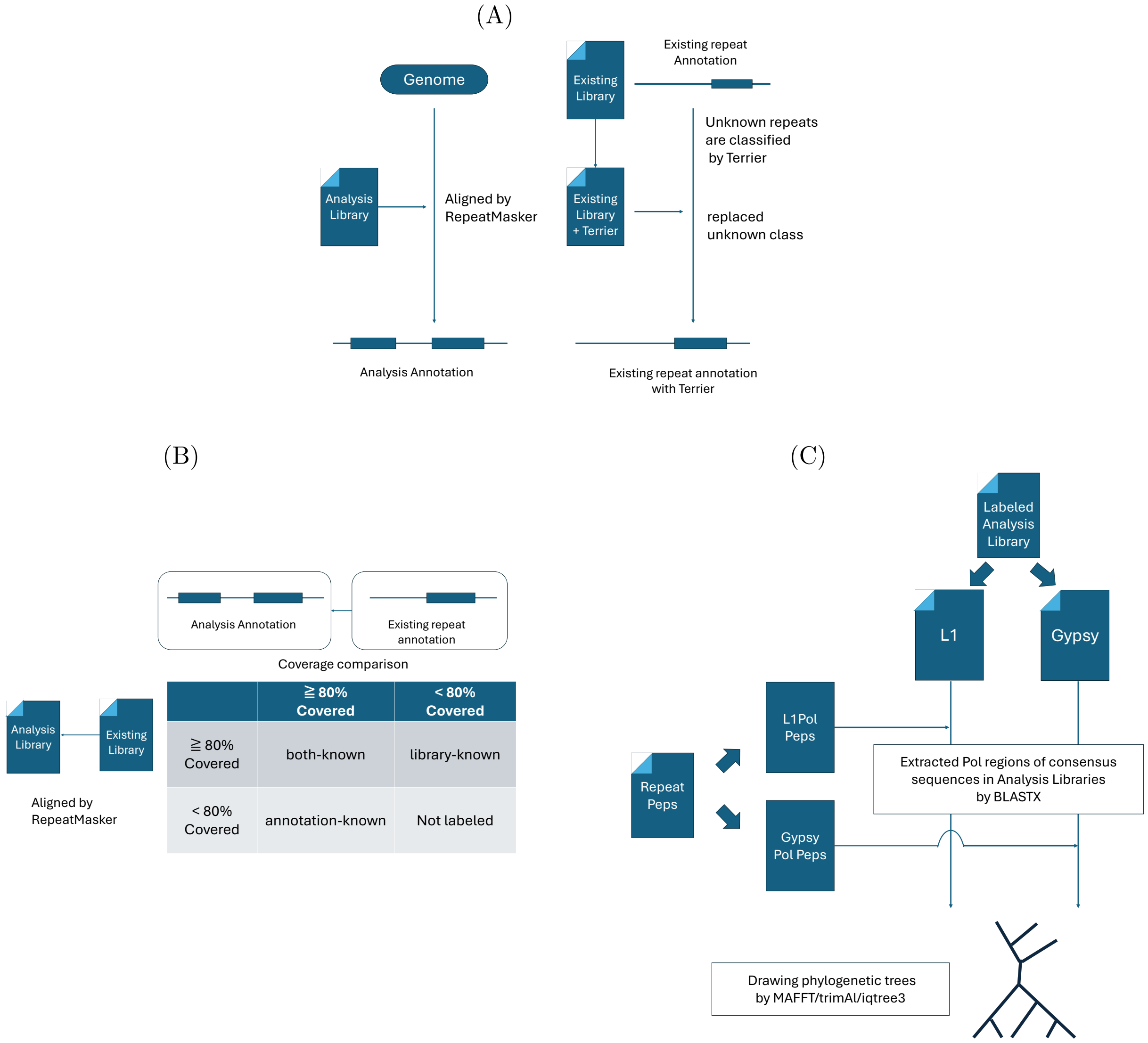


Figure S3: Workflow for generating labeled consensus sequences and Pol-region phylogenetic trees used in the *Pleurodeles waltl* repeat analysis.
(A) Construction of the Analysis Annotation and the Terrier-reclassified existing repeat annotation. The analysis library was aligned to the genome using RepeatMasker to generate the Analysis Annotation. For the existing repeat annotation, repeats annotated as Unknown or Unclassified were reclassified using Terrier, and their class labels were replaced when Terrier assigned a repeat class.
(B) Labeling of consensus sequences in the analysis library. Each consensus sequence was labeled using two coverage-based criteria. In the annotation-based criterion, the RepeatMasker annotation generated from the analysis library was compared with the Terrier-reclassified existing repeat annotation, and a consensus sequence was considered annotation-known when its analysis annotation was covered by the existing repeat annotation by at least 80%. In the library-based criterion, the consensus sequence in the analysis library was compared with the existing repeat library, and it was considered library-known when at least 80% of the analysis-library consensus sequence was covered by the known library. Consensus sequences satisfying both criteria were labeled as both-known, those satisfying only the library-based criterion were labeled as library-known, those satisfying only the annotation-based criterion were labeled as annotation-known, and those satisfying neither criterion were labeled as both-unknown.
(C) Construction of Pol-region phylogenetic trees from labeled consensus sequences. Consensus sequences in the labeled analysis library were searched against Pol protein sequences in RepeatPeps using BLASTX. When a BLASTX hit covered at least 80% of the corresponding Pol protein sequence, the aligned DNA region in the consensus sequence was extracted as a Pol-region sequence. The extracted Pol-region sequences were aligned using MAFFT, trimmed using trimAl, and used to construct phylogenetic trees using IQ-TREE 3.

#### S3 Supplementary Algorithms

**Algorithm S1:** ExpandSeedQueue_DFS

```
Input: FM-index FM = (C, Occ) over BWT L (length n)
SeedQ: stack (or priority queue) of nodes (P, [l, r))
k: target seed length
f_thr: frequency threshold
Mask: masking structure
Output: either (FoundSeed, SeedQ) or (None, SeedQ)
while SeedQ not empty do
    (P, [l, r)) ← pop(SeedQ);
    if |P| = k then
        if EffectiveFreq([l, r), Mask) ≥ f_thr then
            return ((P, [l, r)), SeedQ) ;                    // a "reached" seed
        else
            continue
    // |P| < k: DFS expansion by one character
    foreach a ∈ {A, C, G, T} do
        l' ← C[a] + Occ(a, l);
        r' ← C[a] + Occ(a, r);
        if l' ≥ r' then
            continue
        if EffectiveFreq([l', r'), Mask) < f_thr then
            continue
        push(SeedQ, (aP, [l', r')));
return (None, SeedQ)
Function EffectiveFreq([l, r), Mask):
    return (r − l) − MaskedCount([l, r), Mask)
```

**Algorithm S2:** RightExtension

**Input:** $\{\mathrm{row}_m\}$ $(m = 1..M)$: BWT rows representing the $M$ seed occurrences
$FM$: supports $\mathrm{LF}(\cdot)$, $\mathrm{FL}(\cdot)$, and $\mathrm{BWTChar}(\cdot) \in \{A, C, G, T\}$
$P$: includes $b$ (band half-width), MAXEXTEND, MINIMPROVEMENT, WHEN_TO_STOP
**Output:** $Q_{t_{\mathrm{best}}}$ (right extension of the consensus)

$W \leftarrow 2b + 1$;
Initialize band windows for each occurrence $S_m$ on BWT using FL/LF;
InitScores($P$) ; `// REPrise scoring state for computing` $A(\cdot)$
$Q_0 \leftarrow \varepsilon$;
$t_{\mathrm{best}} \leftarrow 0$;
$A_{\mathrm{best}} \leftarrow A(Q_0; S_1, \ldots, S_M)$;
**for** $t \leftarrow 0$ **to** MAXEXTEND $- 1$ **do**
  $\tau \leftarrow t \bmod W$;
  `// Eq.(1): evaluate` $A(Q_t \cdot N; S_1, \ldots, S_M)$ `for each` $N$
  **foreach** $N \in \{A, C, G, T\}$ **do**
    $A_N \leftarrow \mathrm{ComputeA}(Q_t \cdot N;\ S_1, \ldots, S_M)$;
    ; `// via banded DP (REPrise)`
  $N^* \leftarrow \arg\max_N A_N$;
  $Q_{t+1} \leftarrow Q_t \cdot N^*$ ; `// Eq.(1): update`
  CommitScores($N^*$);
  EmitConsensusBase($N^*, t$);
  `// best-prefix update (RepeatScout/REPrise-style stopping criterion)`
  **if** $A(Q_{t+1}; S_1, \ldots, S_M) \geq A_{\mathrm{best}} + (t + 1 - t_{\mathrm{best}}) \cdot \mathrm{MINIMPROVEMENT}$ **then**
    $t_{\mathrm{best}} \leftarrow t + 1$;
    $Q_{t_{\mathrm{best}}} \leftarrow (Q_{best})$;
    $A_{\mathrm{best}} \leftarrow A(Q_{t_{\mathrm{best}}}; S_1, \ldots, S_M)$;
  **if** $(t + 1) - t_{\mathrm{best}} \geq$ WHEN_TO_STOP **then**
    **break**;
  `// extend neighborhoods of all` $S_m$ `by one symbol on BWT`
  **for** $m \leftarrow 1$ **to** $M$ **do**
    SlideRightOnBWT($\mathrm{row}_m, \mathrm{window}_m, \tau, W, FM$);
**return** $Q_{t_{\mathrm{best}}}$;

**Algorithm S3:** SlideRightOnBWT

**Input:** $\mathrm{row}_m$: current BWT row for occurrence $S_m$
$\mathrm{window}_m$: circular buffer of width $W$
$t$, $W$, $FM$
**Output:** $\mathrm{row}_m$ and $\mathrm{window}_m$ advanced by one symbol

$\tau \leftarrow t \bmod W$;
`// On the genome this would be` $\mathrm{pos} \leftarrow \mathrm{pos} + 1$. `Here we advance via FL/LF on BWT.`
$\mathrm{row}_m \leftarrow \mathrm{FL\_or\_LF}(\mathrm{row}_m)$;
`// FL for forward strand, LF for reverse`
$\mathrm{window}_m[\tau] \leftarrow \mathrm{row}_m$;
**return**

**Algorithm S4:** MaskBWTSpans

```
Input: FM-index with BWT L and LF(·)
spans: set of pairs (from, to) on BWT rows
Mask = {M_A, M_C, M_G, M_T} with rank supports
MAXSTEPS: safety bound
Output: updated Mask and rank supports
foreach (from, to) ∈ spans do
    if from = −1 or to = −1 then
        continue;
    cur ← from;
    steps ← 0;
    while true do
        c ← L[cur] ;                                  // character at BWT row cur
        if c ∈ {A, C, G, T} then
            occ ← Occ(c, cur) ;                       // rank of c in L[0..cur]
            M_c[occ] ← 1 ;                            // mark as masked
        if cur = to then
            break;
        cur ← LF(cur) ;                               // traverse along the span on BWT
        steps ← steps + 1;
        if steps > MAXSTEPS then
            break ;                                   // safety guard
RebuildRankSupports(Mask) ;                           // update rank supports for M_c
```

**Algorithm S5:** RepeatDetectionPipeline

```
Input: FM-index FM over the genome (supports C, Occ, LF, FL, BWT access)
Params includes k, f_min, MAXEXTEND, WHEN_TO_STOP, MINIMPROVEMENT,
MASK_INTERVAL, MAXSTEPS, BATCH
Output: repeat records Records
Mask ← BuildMask(FM);
SeedQ ← empty stack (or priority queue);
push(SeedQ, (ε, [0, n))) ;                                   // root node
masking_count ← 0;
Records ← ∅;
while SeedQ not empty do
    // -- collect a batch of reached seeds (|P| = k) --
    Batch ← ∅;
    while |Batch| < BATCH do
        (Seed, SeedQ) ← ExpandSeedQueue_DFS(FM, SeedQ, k, f_min/2, Mask);
        if Seed = None then
            break;
        // optional: refine frequency at length k
        if EffectiveFreq(Seed.interval, Mask) ≥ f_min then
            add Seed to Batch;
    if Batch is empty then
        break;
    // -- extend seeds (right + left symmetric) --
    (SpansF, SpansR, NewRecords) ← ExtendBatch(FM, Batch, Mask, Params);
    ;    // Right: Algorithm S2 on BWT (LF/FL); Left: symmetric (swap LF and FL)
    append NewRecords to Records;
    // -- masking on BWT --
    MaskBWTSpans(FM, SpansF, Mask, MAXSTEPS);
    MaskBWTSpans(FM, SpansR, Mask, MAXSTEPS);
    masking_count ← masking_count + 1;
    if masking_count mod MASK_INTERVAL = 0 then
        RebuildRankSupports(Mask);
return Records;
```

### S4 Supplementary Results

Table S5: Base-pair-level evaluation counts and derived metrics used for the rice benchmark. TP, FP, and FN indicate true-positive, false-positive, and false-negative base pairs, respectively.

| Tool / Library | TP | FP | FN | Sensitivity | Precision | FDR | F1 |
|---|---|---|---|---|---|---|---|
| BWR-finder(TRs masked) | 15,295,651 | 123,461,421 | 46,566,520 | 0.2473 | 0.1102 | 0.8898 | 0.1525 |
| BWR-finder (TRs not masked) | 27,076,658 | 93,447,861 | 35,398,475 | 0.4334 | 0.2247 | 0.7753 | 0.2959 |
| RepeatModeler2 | 18,664,062 | 54,603,752 | 43,045,696 | 0.3024 | 0.2547 | 0.7453 | 0.2766 |
| HiTE | 12,246,661 | 18,366,146 | 49,647,798 | 0.1979 | 0.4001 | 0.5999 | 0.2648 |
| REPrise | 23,263,822 | 69,003,387 | 40,039,103 | 0.3675 | 0.2521 | 0.7479 | 0.2991 |

Table S6: Base-pair-level evaluation counts and derived metrics for the T2T-CHM13 benchmark. TP, FP, and FN indicate true-positive, false-positive, and false-negative base pairs, respectively. These values were used to generate the corresponding benchmark plot in the main text.

| Tool / Library | TP | FP | FN | Sensitivity | Precision | FDR | F1 |
|---|---|---|---|---|---|---|---|
| BWR-finder (TRs masked) | 94,333,818 | 2,943,613,802 | 346,951,676 | 0.2138 | 0.0311 | 0.9689 | 0.0542 |
| BWR-finder (TRs not masked) | 42,613,247 | 3,549,875,838 | 401,143,904 | 0.0960 | 0.0119 | 0.9881 | 0.0211 |
| RepeatModeler2 | 77,420,851 | 128,254,212 | 368,374,683 | 0.1737 | 0.3764 | 0.6236 | 0.2377 |
| HiTE | 367,263,814 | 521,904,014 | 126,046,969 | 0.7445 | 0.4130 | 0.5870 | 0.5313 |
| REPrise | 30,065,560 | 2,047,032,645 | 414,220,415 | 0.0677 | 0.0145 | 0.9855 | 0.0238 |

Table S7: Computational cost of repeat annotation using the curated library and summary statistics of the repeat library before and after TEstrainer-based curation.

| Category | Item | Value |
|---|---|---|
| RepeatMasker using polished library | Wall-clock time | 413:54:45 |
| RepeatMasker using polished library | Peak memory | 117.3 GB |
| TEstrainer polishing | Number of jobs | 103 |
| TEstrainer polishing | Maximum wall-clock time per job | 32:34:43 |
| TEstrainer polishing | Maximum peak memory per job | 182.1 GB |
| Library before polishing | Number of sequences | 102,578 |
| Library before polishing | Total length | 98.25 Mb |
| Library before polishing | Mean length | 957.8 bp |
| Library before polishing | Median length | 441 bp |
| Library before polishing | N50 | 2,005 bp |
| Library after polishing | Number of sequences | 61,084 |
| Library after polishing | Total length | 167.97 Mb |
| Library after polishing | Mean length | 2,749.8 bp |
| Library after polishing | Median length | 2,346 bp |
| Library after polishing | N50 | 2,824 bp |

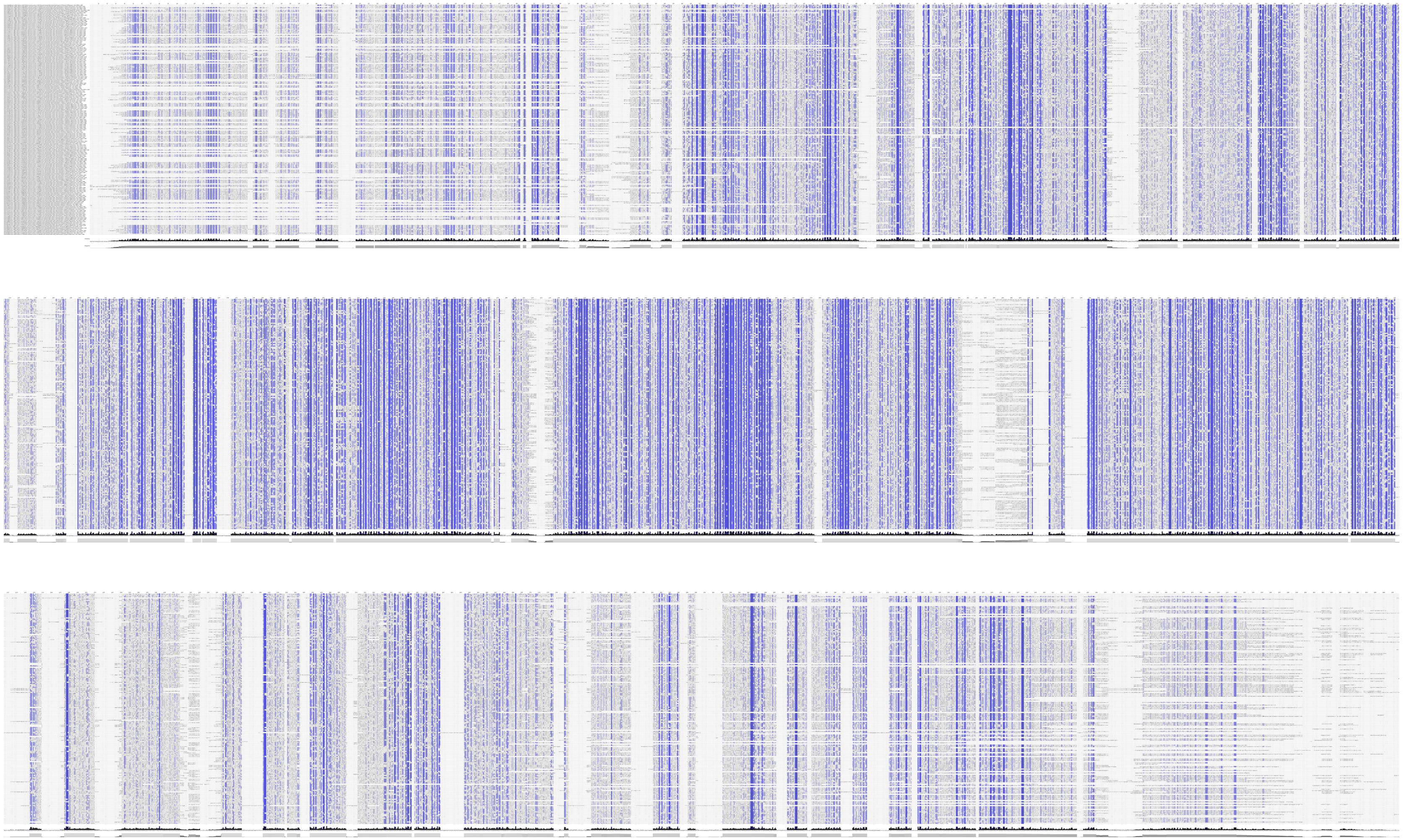

Figure S4: Multiple sequence alignments of the LINE/L1 Pol consensus sequences visualized using Jalview. High identity sites are highlighted according to the coloring scheme in Jalview.

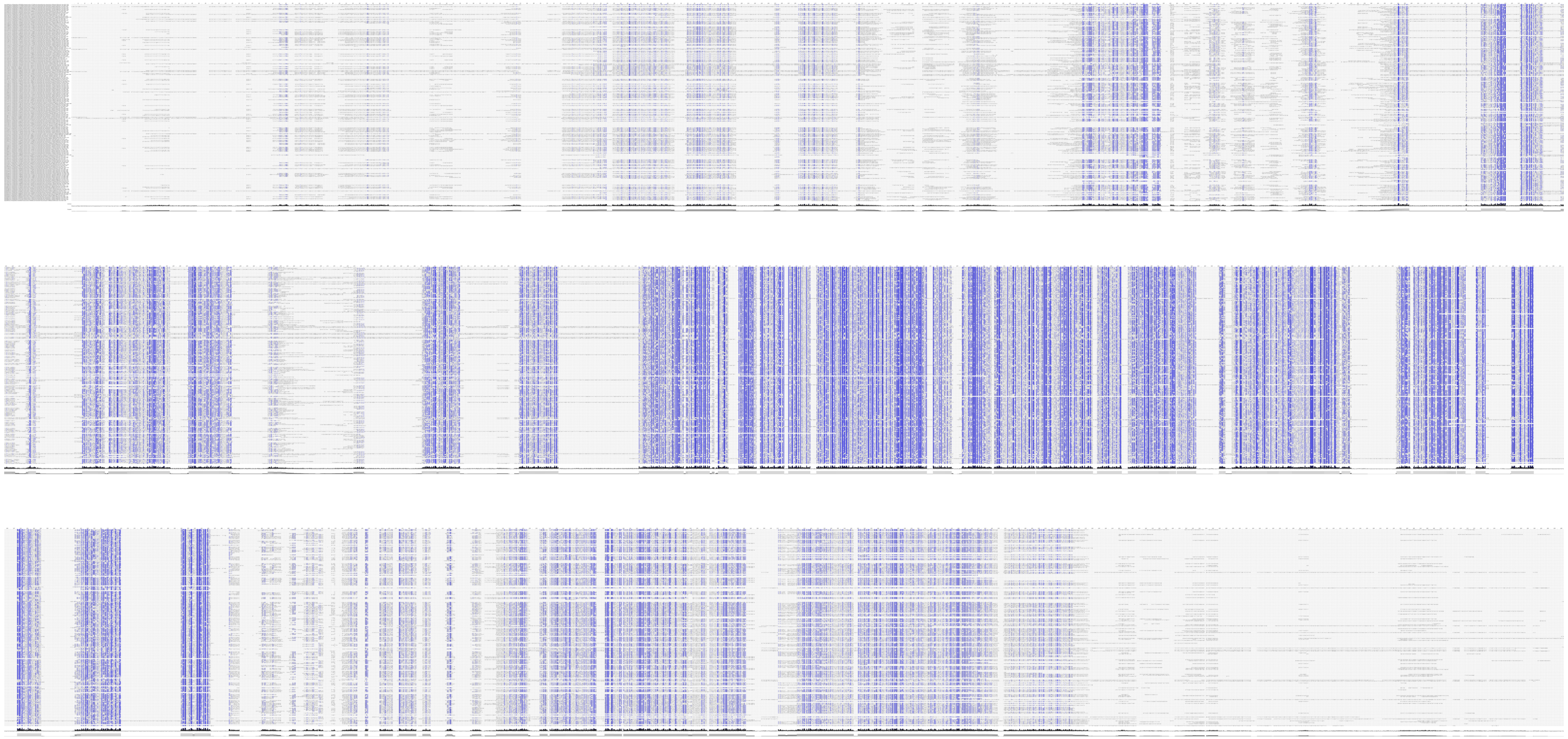

Figure S5: Multiple sequence alignments of the LTR/Gypsy Pol consensus sequences visualized using Jalview. High identity sites are highlighted according to the coloring scheme in Jalview.
